# Functional pocketomes and complete allosteric maps of five protein kinases

**DOI:** 10.64898/2026.08.04.742685

**Authors:** Carla Folgado, Antoni Beltran, Ben Lehner

## Abstract

Protein kinases are central to regulation, dysregulated in many diseases, and the targets of a hundred clinically-approved drugs. Structural conservation of kinase active sites makes the development of selective inhibitors challenging. Targeting allosteric sites can increase selectivity, overcome resistance mutations, and activate kinases. However, most kinases have no known allosteric sites. Moreover, the conservation of allosteric regulation in proteins with the same structural fold is unclear. Here, we address this question by using >390,000 measurements of mutational effects on activity and abundance to construct complete comparative energetic and allosteric maps of five human protein kinase domains: SRC, FGR, JNK2/MAPK9, ZAK/MAP3K20, and TSSK2. For inhibition, all five kinases have distance-dependent allosteric regulation. However allostery is anisotropic, and each protein has a distinct allosteric architecture and surface to therapeutically target. A set of inhibitory allosteric pockets close to the active site is conserved in all five proteins, but more distal allosteric pockets are protein-specific. These include pockets that are activatory in one kinase and inhibitory in another. Differences in allosteric regulation are particularly striking for activation, where the allosteric maps are highly diverged. Our results show that proteins with the same structural fold have different energetic architectures and allosteric pockets. Mapping the ‘functional pocketomes’ of kinases and other clinically-important targets should empower the development of more selective therapeutics.

## Introduction

The human genome encodes more than 500 protein kinases, proteins that catalyze the transfer of a phosphate group from adenosine triphosphate (ATP) to specific serine, threonine, or tyrosine residues of target proteins to regulate their activity. Kinases are central to biological control and to nearly all signal transduction pathways, regulating processes as diverse as metabolism, growth, cell division, transcription, DNA repair, and neuronal communication^1^. Kinases are frequently dysregulated and causally mutated in human diseases, particularly in cancer^2–4^. As a result, kinases are amongst the most important classes of human drug targets^5^, with nearly 100 kinase-targeting drugs now approved for clinical use^6^. These drugs collectively target ∼50 different kinases^7^.

Protein kinases are classic allosteric proteins, with their activities physiologically controlled by molecular interactions and modifications outside of their active sites^8^. Despite this, nearly all kinase drugs bind the highly-conserved ATP-binding pocket. As for most enzyme active sites, this orthosteric pocket is highly structurally conserved in the kinase family, which makes it very challenging to selectively inhibit one kinase without also inhibiting many others^9–12^. For example, dasatinib, used to treat chronic myeloid leukaemia, binds at least 52 kinases at high affinity (Kd < 100 nM), and sorafenib, approved for renal and hepatocellular carcinoma, binds at least 16^13^. This lack of specificity is a frequent cause of toxicity and clinical failure^14^. Moreover, resistance mutations frequently arise in the active sites of kinases, rendering active-site targeting compounds ineffective^15^. One approach to increase specificity is to target functional secondary sites. Allosteric drugs binding secondary sites outside of the active site have been developed for multiple kinases^16^, including asciminib, that targets the ABL1 myristate binding site^17,18^, the MEK1/2 inhibitors trametinib and selumetinib, that bind a pocket located in the vicinity of the ATP-binding site and helix αC^19^, and deucravacitinib^20^, that binds the pseudokinase domain of TYK1 for allosteric inhibition. Allosteric drugs can also overcome active site resistance mutations^21^ and small molecules binding secondary sites can activate not just inhibit kinase activity^22,23^.

For the vast majority of kinases — and other proteins — however, the functional secondary sites to target are unknown. At one extreme it could be that proteins with conserved structures also have highly observed energetic architectures, with conserved allosteric networks and the same functional secondary sites and allosteric pockets. At another extreme, despite sharing the same 3D-fold, each protein in a structural family could have a very different energetic architecture, with divergent allosteric networks and non-conserved allosteric surface sites and unique allosteric pockets. It is the question that we address in this manuscript: to what extent are the energetic and allosteric networks and functional secondary sites conserved in proteins with the same structural fold?

Allosteric drugs have nearly all been serendipitously discovered^24^. However, the development of high-throughput methods for allosteric mapping provides an alternative strategy of directly identifying functional sites to target in each protein^25,26^. In this approach, mutations are used as perturbations that introduce 19 different changes in chemistry at each site. Multi-modal selections and model fitting are then used to quantify changes in protein activity beyond those caused by changes in the abundance of the folded protein^25,26^. Applied to the SRC kinase, this approach produced the first comprehensive map of allostery for an enzyme^27^. Allostery in SRC is distance-dependent, with mutations close to the active site much more likely to inhibit enzymatic activity. However, the energetic architecture is asymmetric, with stronger allostery in particular directions within the 3D structure. Moreover, activating mutations in SRC have a very different structural distribution to inactivating mutations. The SRC allosteric map identified allosteric surface pockets targeted by known ligands, but also multiple new functional secondary sites to potentially therapeutically target^27^.

Here we use the approach of generating complete allosteric maps to directly address the question of the extent of conservation of allosteric regulation and allosteric sites in proteins with conserved three dimensional structures. We target five human protein kinases from across the kinase family phylogenetic tree: SRC, FGR, MAPK9, ZAK, and TSSK2. The five kinases also differ in their physiological roles, disease associations and the extent to which they have been successfully drugged. SRC is the prototypical oncogenic tyrosine kinase, with hyperactivation implicated in the progression and invasion of breast, colorectal and other solid tumours^28^. Although its ATP-competitive inhibitors dasatinib and bosutinib are clinically approved, these are primarily used to treat BCR-ABL-driven leukaemias and are not selective SRC inhibitors^28,29^. FGR is a myeloid-restricted SRC-family kinase active in neutrophils, monocytes and macrophages, where it participates in integrin-dependent activation and inflammation^30,31^, and regulates mitochondrial complex II activity to support metabolic adaptation^32^. FGR is a target in inflammatory metabolic disease, and in acute myeloid leukaemia, where a selective inhibitor suppresses leukaemic cell growth^33^, though no FGR-directed agent has entered the clinic. JNK2 (MAPK9) is a stress-activated serine/threonine kinase of the JNK family that phosphorylates AP-1 transcription factors such as c-Jun and ATF2, with context-dependent roles in stress signalling, cancer and immunity^34,35^. The JNK pathway has been pursued therapeutically^36^, but no isoform-selective JNK2 inhibitor has been approved^37^. ZAK (MAP3K20) is a stress-activated MAP3K that signals through the JNK and p38 pathways to trigger cell-cycle arrest and programmed cell death. Its two splice isoforms act in distinct biological contexts: ZAKα mediates the ribotoxic stress response to stalled ribosomes and ribosome-targeting toxins in most tissues^38^, whereas ZAKβ is the isoform expressed in skeletal muscle, where it is activated by muscle contraction and mechanical compression and is required for the adaptive response to sustained mechanical load^39^. Loss of ZAK function causes a congenital myopathy with progressive muscle weakness and defective Filamin C turnover, and it remains a genetically-validated but largely undrugged target^40,41^. Finally, TSSK2 is a testis-specific serine/threonine kinase required for sperm maturation and male fertility, and is of interest as a candidate target for non-hormonal male contraception^42^, though no inhibitor has reached the clinic. Together, these proteins span the kinome: two SRC-family tyrosine kinases and three serine/threonine kinases from distinct groups, and a broad range of physiological contexts, disease associations and stages of therapeutic development, from approved but non-selective inhibitors to targets that have never been drugged.

The five allosteric maps that we present here provide, to the best of our knowledge, the first opportunity to comprehensively compare the energetic and allosteric architectures of homologous proteins from an enzyme family. Whilst certain aspects of the allosteric maps are highly conserved, including the principle of anisotropic distance-dependent allosteric decay for inhibition, other features are protein-specific. Comparing potentially druggable surface pockets identifies a set of pockets close to the active site - including the DFG and MT3 pockets - that is allosteric in all five kinases. However, further from the active site, each kinase has a different set of allosteric pockets and functional secondary sites. This divergence is particularly strong for protein activation. Each kinase therefore has a distinct energetic and allosteric architecture and a distinct set of functional surface pockets to potentially therapeutically target. More generally, our results show that proteins with the same structural fold can have diverged energetic and allosteric architectures. Mapping the ‘functional pocketomes’ of kinases and other proteins should help unlock the development of more selective modulators.

## Results

### Charting complete allosteric maps for five human kinases

To compare the energetic and allosteric structures of different proteins that share a conserved structural fold we used an experimental platform that we refer to as KINASE-MAPS (Kinase-Mutational Allosteric Propagation Scan) to quantify the effects of all possible amino acid (aa) substitutions on both the activity and folded abundance of five human protein kinases^27^. KINASE-MAPS has four steps: First, we construct a mutant library for each kinase that, importantly, consists of combinatorial mutants (here, double mutants) to constrain model-fitting (Figure 1F). Second, we use selection-sequencing experiments (Figure S1H) to quantify the kinase activity of every library variant (Figure 1E). Third, we quantify the folded protein abundance of every library variant (Figure 1E). Fourth, we fit an energy model^43^ to the combined data to infer the change in fold stability and activity for each single aa variant (Figures 1B and 1C).

**Figure 1.**
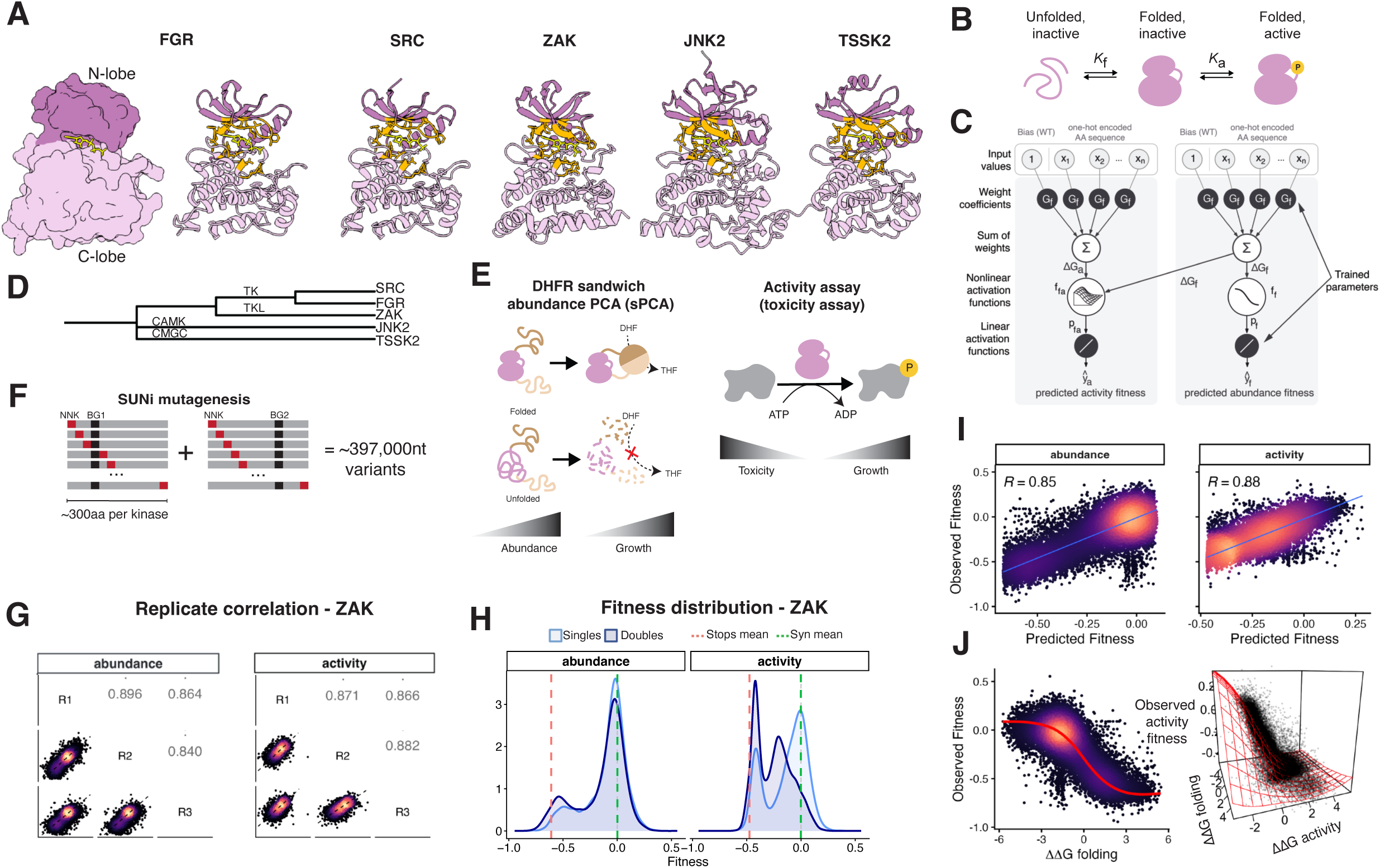
Charting comparative energetic and allosteric maps for five protein kinases. (A) AlphaFold predicted structures of the kinase domains studied (FGR, SRC, ZAK, JNK2, TSSK2). N-lobes are shown in dark pink, and C-lobes in light pink. ATP-contacting residues are shown in orange. ATP is shown as yellow sticks. (B) Three-state model of kinase folding and activation, with the states linked by the folding (Kf) and activation (Ka) equilibrium constants. (C) Architecture of the MoCHI model. Additive folding (ΔGf) and activity (ΔGa) free energies, inferred from the amino acid sequence, are transformed into predicted abundance and activity fitness. (D) Dendrogram of the five kinases and their kinase family classification (TK, TKL, CAMK, CMGC). (E) Overview of the two selection assays. In the DHFR abundance protein-fragment complementation assay (sPCA), yeast growth increases with the amount of folded protein (DHF, dihydrofolate; THF, tetrahydrofolate). In the activity (toxicity) assay, yeast growth decreases with kinase activity. (F) SUNi mutagenesis strategy. We introduced NNK codons encoding all single amino acid substitutions across ∼300 residues per kinase, in 8 genetic backgrounds (in two residues, BG1 and BG2, with four mutations each). (G) Fitness correlation between the three replicates (R1 to R3) of the abundance and activity assays for ZAK (Pearson’s R). (H) Fitness distributions of single and double mutants of ZAK for the abundance and activity assays; dashed lines mark the mean of stop codons (red) and synonymous variants (green). (I) Observed against MoCHI-predicted fitness for ZAK in the abundance and activity assays (Pearson’s r). (J) Observed abundance fitness of ZAK as a function of the inferred free energy of folding (ΔΔGf, left; red, fitted model) and as a function of both ΔΔGf and the inferred free energy of activity (ΔΔGa, right).

As representative kinases we targeted two SRC-family tyrosine kinases (SRC, FGR) and three serine/threonine kinases: a CMGC kinase (JNK2/MAPK9), a TKL kinase (ZAK/MAP3K20), and a CAMK kinase (TSSK2), providing broad sampling of the protein kinase phylogenetic tree (Figure 1D). Data for four kinases were generated for this study, while the SRC data was previously published^27^. The five proteins vary in length from 358 (TSSK2) to 536 aa (SRC). SRC and FGR have N-terminal SH3 and SH2 domains followed by a kinase domain and a C-terminal regulatory tail, ZAK isoform B (ZAKβ) comprises an N-terminal kinase domain followed by a SAM domain, and JNK2 and TSSK2 each comprise a single kinase domain followed by a disordered region (Figure S1I).

We used SUNi mutagenesis^44^ to generate shallow double aa mutant libraries covering the entire 269 to 357 aa kinase domain (Figure 1A) of each protein, expressing them in their full-length format. These libraries encode all possible single kinase domain aa substitutions within the full-length kinase construct, in combination with eight ‘background’ aa changes (four background variants at two different sites) (Figure 1F). The library for each kinase therefore contains an average of 78,579 nucleotide (nt) variants encoding 50,510 aa changes and the total design contains 396,800 nt variants, encoding 248,000 distinct aa sequences.

We quantified the activity of each double mutant variant of each kinase using a highly-validated cell-based toxicity assay^27,45–47^ (Figure 1E). For all five proteins we validated that toxicity was activity-dependent using active site mutations (Figures S1A and S1D). The activity selections were well correlated across three independent experimental replicates (median r=0.82 for single mutants; median r=0.77 in the full dataset including higher order variants, Figures 1G and S1B). Many changes in protein activity are caused by changes in the concentration of folded protein^27,43,48^, so we also quantified the folded concentration of each protein in the same cellular context using a second growth-based selection, sandwich protein-fragment complementation assay (sPCA)^27,49^ (Figure 1E). Abundance selections were also highly reproducible (median r=0.84 for single mutants; median r=0.78 full dataset, Figures 1I and S1B) and all datasets had good separation between stop and synonymous variants for both activity and abundance (Figures 1H and S1C). After quality control, we quantified the activity of 163,185 aa variants and the folded abundance of 231,716 variants (Table S1, S1F and S1G), with each aa substitution represented in a median of 5 double mutants (Figures S1E).

In the final step we fit an energy model to the data for each kinase. The goal is to quantify changes in kinase activity not accounted for by changes in the folded abundance of each protein. We use a simple phenomenological model in which each protein can exist in three states — unfolded, folded inactive, and folded active (Figure 1B) — with the population distributions across these states governed by two free energy terms, the Gibbs free energy of folding (ΔGf) and an activity energy (ΔGa) (Figures 1C and S2C). The activity energy captures all changes in kinase activity not accounted for by changes in folded protein abundance, including shifts in the equilibrium between active and inactive states and changes in the catalytic parameters^27^. Mutational effects on folding (ΔΔGf) and activity (ΔΔGa) are assumed to combine additively in double mutants. Evaluated by ten-fold cross validation, the model provides excellent prediction of the abundance (median r=0.78, with attenuation correction r = 0.85) and activity (median r=0.81, with attenuation correction r = 0.90) of double mutants (Figures S2A and S2B). In total, the resulting dataset quantifies 50,396 changes in free energy across the five kinases: 26,032 changes in fold stability (ΔΔGf) and 24,711 changes in activity (ΔΔGa) (Figures S3A, S3B, and Table S2).

### Conservation and divergence of mutational effects on fold stability

We first compared mutational effects on fold stability (ΔΔGf) across the five kinase domains (Figures 2, S4A and S4B). Previous comparisons in small domains (<85 aa) have suggested mutational effects are highly conserved in protein families^50^. However, the kinase domain is a large and multi-lobed fold. Between 881 and 2280 mutations destabilised each kinase (ΔΔGf > 0.2 kcal/mol, one-sided z-test, Benjamini-Hochberg (BH) FDR<0.1). As expected, destabilising mutations are enriched in the protein core in all five kinases (OR = 4.0, p < 0.05, one-sided Fisher’s exact test (FET)), with the level of enrichment ranging from OR = 2.0 (TSSK2) to OR = 11.5 (ZAK), all p < 0.05 (Figure S4C). Destabilising mutations are also enriched in the catalytic loop (CL) and in the αF and αE helices (all in 4 of 5 kinases, Figure S4G).

**Figure 2.**
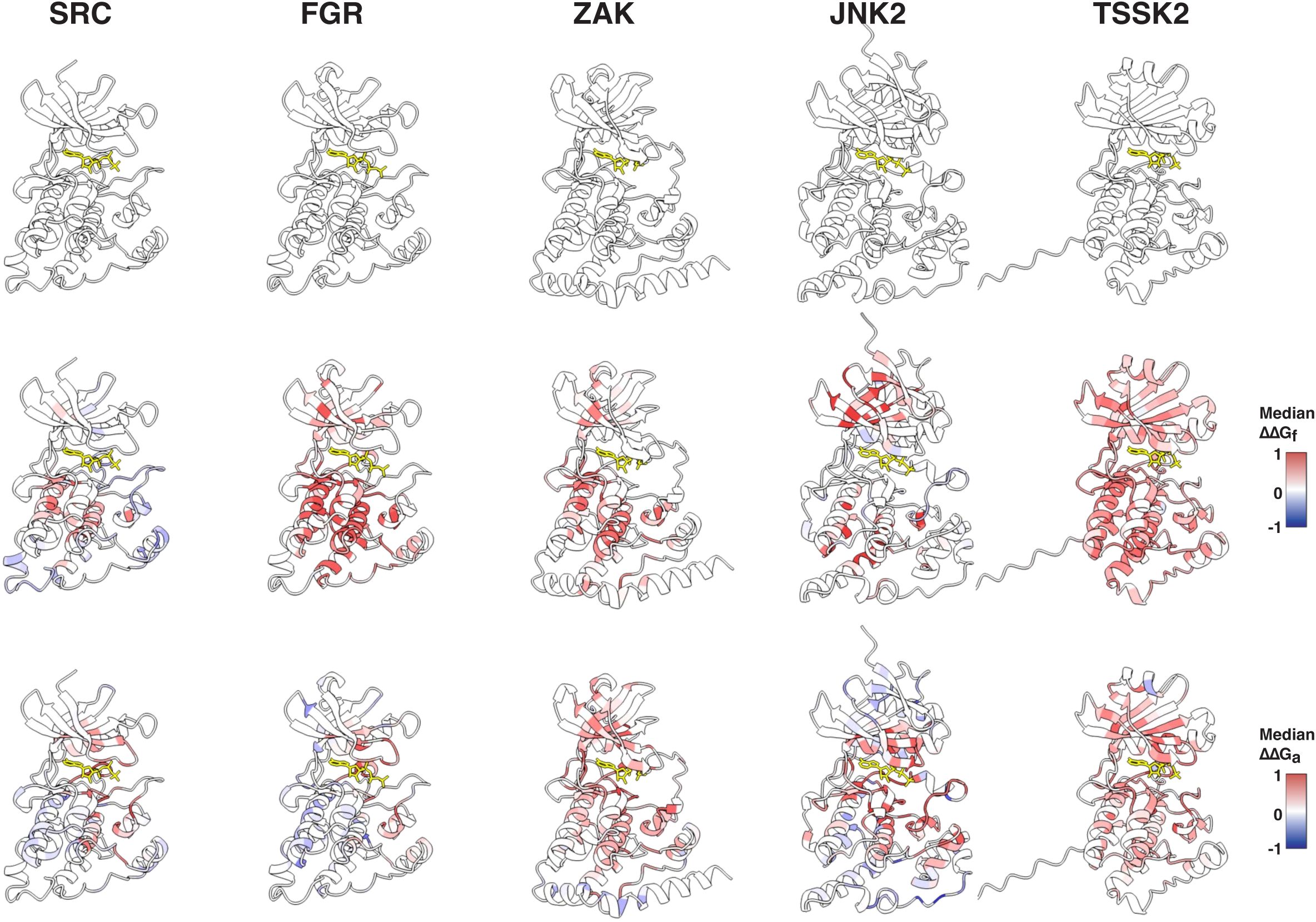
Folding and activity energetic landscapes. AlphaFold3 structures of the kinase domains of SRC, FGR, ZAK, JNK2 and TSSK2, shown uncoloured (top row) and coloured by the per-position median ΔΔGf (middle row) or ΔΔGa (bottom row). ATP is shown as yellow sticks.

Interestingly, however, the importance of different structural regions for fold stability varies across the five proteins. Whereas in SRC, FGR and ZAK destabilizing mutations are enriched in the C-lobe (OR = 1.7, 2.3 and 1.7, respectively, FET, p < 0.05), in JNK2 they are enriched in the N-lobe (OR = 2.2, FET, p < 0.05), and in TSSK2 they are similarly frequent in both lobes (63.7% vs 65.0%, p = 0.48) (Figure S4F). Mutational effects on stability are well-correlated between the two most related kinases, SRC and FGR (Pearson r = 0.77), and also quite well correlated with the next most-related kinase, ZAK (r = 0.66 and r = 0.68 for SRC and FGR, respectively) (Figure S4E). Mutational effects on stability are more divergent in JNK2 and TSSK2 (SRC r = 0.40 and r = 0.39, respectively, median with all kinases r = 0.39) (Figure S4E). Considering individual secondary structures and regions (Figure S4D), mutational effects on stability in the β2 strand and the αH helix are most correlated (both with median r = 0.62 across 10 kinase pairs), whereas mutations in β4, αG and αC are most divergent (median r = 0.19, 0.20 and 0.22, respectively). Even within the αF helix, which is the most buried secondary structure element (mean relative solvent accessibility (rSASA) = 0.11) and the one where mutations are most destabilising (median ΔΔGf = 0.39 kcal/mol), mutational effects vary quite extensively (Figure S4I), with mutations more detrimental for stability in TSSK2 and FGR in the C-terminal 12 residues of αF (Figures S4H, S4I and S4J 2B).

Thus, mutational effects on stability vary across different regions in different kinases, including in buried secondary structures.

### Stabilizing mutations

Across the five kinases, 1,578 mutations increase protein abundance (ΔΔGf < -0.2 kcal/mol, one-sided z-test, FDR < 0.1), ranging from 75 (TSSK2) to 978 (SRC). The activation loop (AL) is the only region enriched for stabilizing substitutions in multiple kinases (OR = 7.3 (SRC), 3.2 (JNK2), 2.2 (FGR), 1.7 (ZAK) (FDR < 0.1, FET, for SRC, JNK2 and FGR) (Figure S4G). In TSSK2, in contrast, the AL is not enriched for stabilizing substitutions (OR = 0.64, FET, FDR = 0.65). Four of the five kinases also have protein-specific regions enriched for stabilising mutations: αG and the F-loop linker in SRC, αD in JNK2, αC and β1 in FGR, and β2 and β5 in TSSK2 (Figure S4G, all FET FDR < 0.1). No regions are enriched for stabilizing mutations in ZAK (FDR < 0.1). The landscape for stabilising mutations thus varies quite extensively across five enzymes in the same protein family.

### Active sites

Our data provide the first complete comparative analysis of the effects of mutations on protein kinase activity independent of their effects on protein abundance. Across the five kinases we identify a total of 3,996 inactivating mutations (ΔΔGa > 0.2 kcal/mol, one-sided z-test, FDR < 0.1) and a total of 1,087 activating mutations (ΔΔGa < −0.2 kcal/mol, one-sided z-test, FDR < 0.1, Figure 2). The number of inactivating mutations ranges from 352 (FGR) to 1,216 (JNK2) and the number of activating mutations ranges from 19 (TSSK2) to 474 (JNK2). We first consider the effects of mutations in the active site of each kinase, defined as residues contacting ATP in any of the five kinases (22 residues), the magnesium-positioning loop (3 residues) and the substrate-positioning loop (10 residues), totalling 35 aligned sites with 2,954 energetic measurements across the five kinases. Active sites are enriched for inactivating mutations in all five proteins (n = 1,379, pooled OR = 6.4, OR = 4.1 to 11.2 per kinase, FET, all FDR < 0.1) and depleted for activating mutations in three of five (n = 47; pooled OR = 0.32; FGR, JNK2 and ZAK; OR = 0 to 0.42, FET, FDR<0.1; Figure S5B). Mutations in the magnesium-positioning loop are highly detrimental in all five proteins (median of mutations ΔΔGa = 0.86 kcal/mol), as are mutations in the catalytic-loop ATP contacts (alignment positions 161, 163, 165, 166 and 168, median ΔΔGa = 0.65 kcal/mol), the N-terminal half of the substrate-positioning segment of the activation loop (alignment positions 202-206; median of mutations ΔΔGa = 0.62 kcal/mol), and the glycine-rich loop (alignment positions 34-39,median of mutations ΔΔGa = 0.53 kcal/mol), (Figures 3A, 3B and S5A).

**Figure 3.**
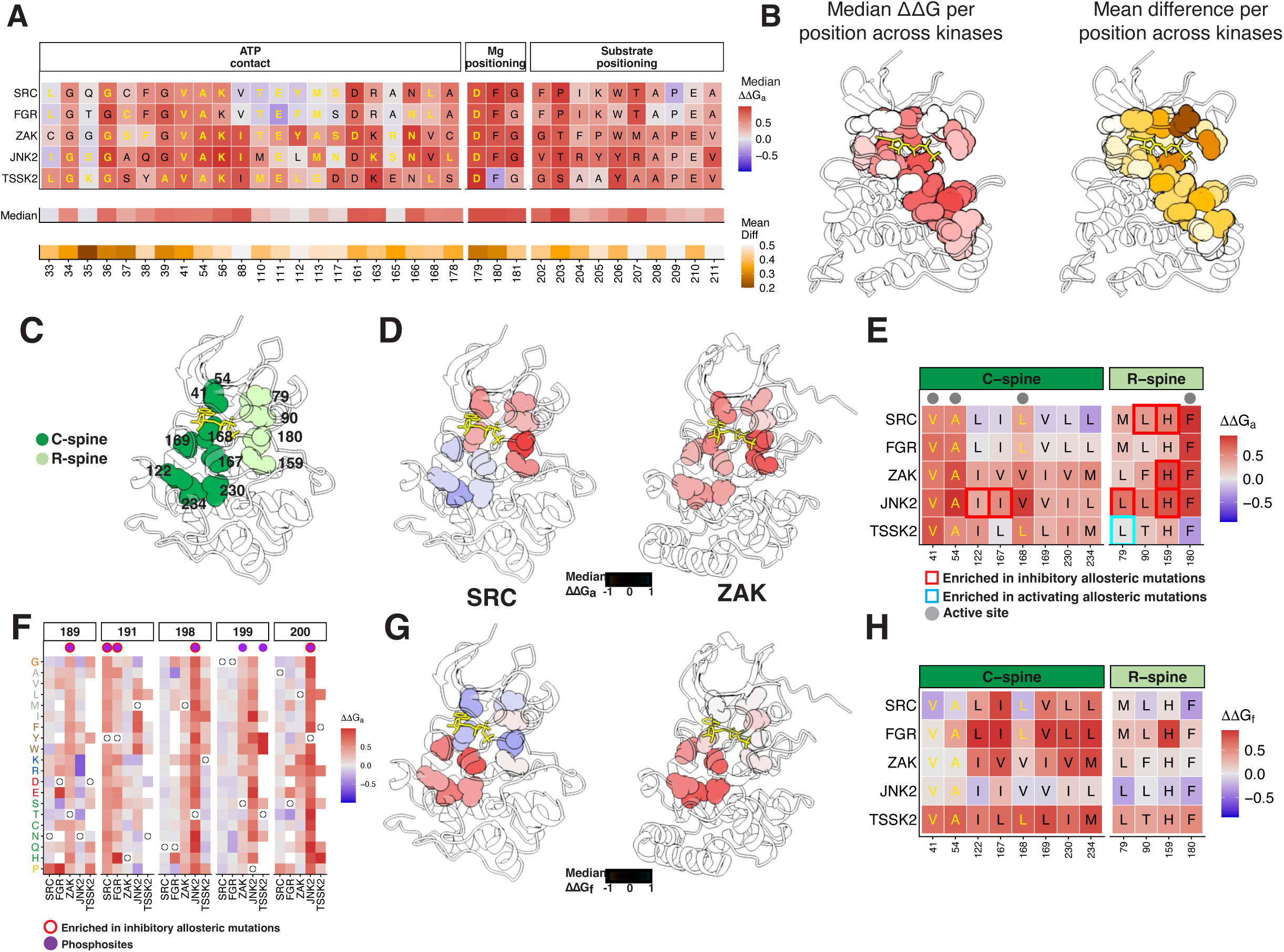
Mutational effects at the active site and known regulatory sites. (A) Top, median ΔΔGa per position at the active-site residues of each kinase, grouped by function (ATP contact, Mg-positioning, substrate-positioning), mapped into aligned positions. Letters are the wild-type residue (ATP contacts in yellow). Middle, median ΔΔGa per position across kinases. Bottom, cross-kinase divergence (mean pairwise absolute difference in ΔΔGa across kinases). (B) Active-site positions mapped onto the SRC structure as spheres, coloured by the median ΔΔGa across kinases (left) and by the cross-kinase divergence (right). ATP is shown as yellow sticks. (C) Catalytic spine (C-spine, dark green) and regulatory spine (R-spine, light green) positions on the structure, labelled with aligned position numbers. (D) C-spine and R-spine positions on the SRC and ZAK structures, coloured by per-position median ΔΔGa. (E) Median ΔΔGa at C-spine and R-spine positions for each kinase. Letters are wild-type residue, yellow if ATP contact. (F) ΔΔGa off all substitutions across the five kinases (rows, amino acids) at activation-loop positions that are phosphosites in at least one kinase. White circles mark the wild-type residue. Annotated phosphosites in each kinase are marked by purple circles. (G) C-spine and R-spine positions on the SRC and ZAK structures, coloured by median ΔΔGf. (H) Median ΔΔGf at C-spine and R-spine positions for each kinase.

In contrast, mutations in the six residues that contact the adenosine portion of ATP have weaker effects (alignment positions 88, 110-113, 117; median ΔΔGa = 0.19 kcal/mol) that also vary more across the five proteins (mean pairwise difference between kinases in adenosine contacts = 0.47 kcal/mol, vs 0.36 kcal/mol in the rest of the ATP-contact residues) (Figures 3A and 3B). Mutations in the conserved glutamate at position 111 are enriched for activating mutations in both tyrosine kinases (SRC 3/19, OR = 10.7; FGR 5/16, OR = 15.5; FET, both p < 0.05) but not in JNK2, ZAK and TSSK2 (0/19, 0/19 and 0/12). E111 is the residue immediately C-terminal to the gatekeeper (T110), where mutations are also activating in SRC and FGR. The active site thus divides into two parts: a catalytic core (the magnesium-positioning loop, the catalytic-loop ATP contacts, the glycine-rich loop and the N-terminal activation loop) that is intolerant to mutation in all five kinases, and an adenosine-binding region where effects diverge, with a subset of mutations activating the two tyrosine kinases but not the three serine/threonine kinases.

### Known regulatory sites

We next consider allosteric mutations - mutations outside of the active site that modulate activity. In total we identified 2,617 inhibitory allosteric mutations (SRC 375, FGR 203, ZAK 810, JNK2 842, TSSK2 387) and 1,040 activating allosteric mutations (SRC 150, FGR 309, ZAK 100, JNK2 463, TSSK2 18) (one-sided z-test, FDR < 0.1).

Kinase domains contain two structurally distinct hydrophobic ‘spines’ that link the N- and C-lobes and create a rigid hydrophobic core aligning the catalytic residues for efficient phosphoryl transfer. The Regulatory (R)-spine is formed only in the active state, and the Catalytic (C)-spine is completed by the adenine ring of ATP^51,52^ (Figure 3C). Mutations in the four R-spine residues (79, 90, 159 and 180) have little effect on fold stability (median ΔΔGf = 0.08 kcal/mol, Figures 3G and 3H) but frequently inhibit activity (median ΔΔGa = 0.53 kcal/mol, Figures 3D and 3E). In the C-spine, mutations in the three ATP-contacting positions (41, 54 and 168) have the largest effects on activity (median ΔΔGa = 0.58 kcal/mol), with two further positions in JNK2 (122 and 167, both isoleucine) also enriched for inhibitory mutations (FET, FDR < 0.1, median ΔΔGa = 0.33 and 0.50 kcal/mol) (Figures 3D, 3E, 3G and 3H). In contrast, mutations in the remaining C-spine positions have smaller effects on activity (median ΔΔGa = 0.18 kcal/mol) and instead affect stability (median ΔΔGf = 0.67 kcal/mol). The αF helix, the central helix of the C-lobe that anchors both the R and C-spines, is enriched in allosteric mutations in three out of five kinases (SRC, ZAK and JNK2; OR = 1.4 - 1.9, FDR < 0.1) (Figures S6D and S6F).

The αC-helix acts as a conserved allosteric lever, swinging between active and inactive orientations to gate catalysis^52,53^. The conserved αC-helix glutamate (alignment position 75) forms the regulatory salt bridge with the β3 lysine and is the only position enriched for inhibitory allosteric mutations in all five kinases (OR = 8.3 - 93.7, FET, FDR < 0.1 each, median ΔΔGa = 0.57 to 0.95 kcal/mol). The rest of the αC helix is enriched in allosteric mutations in FGR, JNK2 and TSSK2 (OR = 1.7 - 2.5, FET, FDR < 0.1), but not in SRC (median ΔΔGa = 0.07 kcal/mol) or ZAK (median ΔΔGa = 0.2 kcal/mol) (Figure S7A).

Phosphorylation of the activation loop triggers conformational changes for kinase activation^54^. The activation loop is enriched for inhibitory mutations in three of the five kinases (JNK2 OR = 4.21, FGR OR = 1.86, SRC OR = 1.65; FET, FDR < 0.1) but not in ZAK or TSSK2 (OR = 0.93 and 0.87, FDR = 0.75 and 0.62) (Figure S6D). The activation loop contains seven annotated phosphosites across the five kinases and 104 of 120 substitutions at these sites are inactivating and none are activating (Figure 3F). Serine to threonine substitutions that retain a potentially phosphorylatable hydroxyl are inactivating at three sites (JNK2 198, ZAK 189 and TSSK2 199; ΔΔGa = 0.19 to 0.69 kcal/mol) but not at ZAK 199 (ΔΔGa = -0.11 kcal/mol). Potentially phosphomimetic aspartate and glutamate mutations are not activating at any activation-loop serine, threonine or tyrosine (ΔΔGa = -0.06 to 1.12 kcal/mol).

SRC and FGR activity is inhibited by binding of their SH3 and SH2 domains onto the side of the kinase domain distal to the active-site cleft, with the SH2 docking against the αE helix at the C-lobe, and the SH3 domain binding to the SH2-kinase linker^55–57^. Across the αE helix (aligned positions 133-151), activating mutations are enriched at 5 of 19 surface positions in SRC (OR = 7.8-118; FET, FDR < 0.1) and 3 of 19 in FGR (OR = 5.8-8.1; FET, FDR < 0.1) (Figures S6D and S7A). Among all the N-lobe positions contacted by the SH3 domain (SRC 46, 47, 50, 51; FGR 7, 46, 47, 50, 51, 83) and by the SH2-kinase linker (SRC 46, 51, 53, 83, 89, 91, 92, 111; FGR 89, 91, 111, 171, 172) only FGR position 46 (SH3 contact),is enriched for activating mutations (OR = 7.4, FET, FDR < 0.1) (Figure S7A).

JNK2 binds its activating kinases (MKK4 and MKK7), substrates and phosphatases through the D-recruitment groove^58^, which is enriched for inactivating mutations at position 172 in the activation loop (OR = 8.3, FET, FDR < 0.1, ΔΔGa = 0.48 kcal/mol) and position 341 in the common-docking site C-terminal to αG (OR = 10.4, FET, FDR < 0.1, ΔΔGa = 0.51 kcal/mol) (Figures S7A). Mutations in the rest of the groove only have small effects (ΔΔGa -0.05 to 0.14 kcal/mol). To our knowledge, ZAK and TSSK2 do not have any additional known regulatory sites in their kinase domains.

### Conserved distance-dependent allosteric decay for kinase inhibition

We next considered the spatial arrangement of inhibitory allosteric mutations across the kinase domains. Mutations close to the active site are much more likely to be allosteric in all five kinases. This distance-dependent allostery is well illustrated by plotting the median ΔΔGa for all inhibitory mutations at each residue (Figure 4B), or the median |ΔΔGa| for all mutations (Figure S6A), against the distance to the active site (minimum heavy-atom distance to the ATP nucleotide or the catalytic aspartate) (Figure 4A). Fitting exponential decay functions quantifies the distance over which the median inhibitory allosteric effect halves (d½) as 9.3 Å across the five proteins, ranging from 6.5 Å in SRC (95% confidence interval (CI) 5.-8.6) to 11.1 Å in ZAK (95% CI 9.0-14.3). SRC has the sharpest decay, and its 95% CI does not overlap that of ZAK (11.1 Å, 95% CI 9.0-14.3) or TSSK2 (10.9 Å, 95% CI 9.2-13.4) (Figure 4C). The confidence intervals of the remaining kinases overlap, indicating similar decay rates. A half-distance of 9.3 Å corresponds to roughly two successive shells of amino-acid contacts (mean spacing 4.5 Å per shell), with the median inhibitory effect of a mutation reducing 28.5% per shell and approximately halving with a distance of two contacts. The conserved distance-dependent decay of allostery^25,26,59–61^ is consistent with measurements of energetic couplings and structural perturbations^62^ and may underlie evolutionary conservation gradients away from active sites^63,64^.

**Figure 4.**
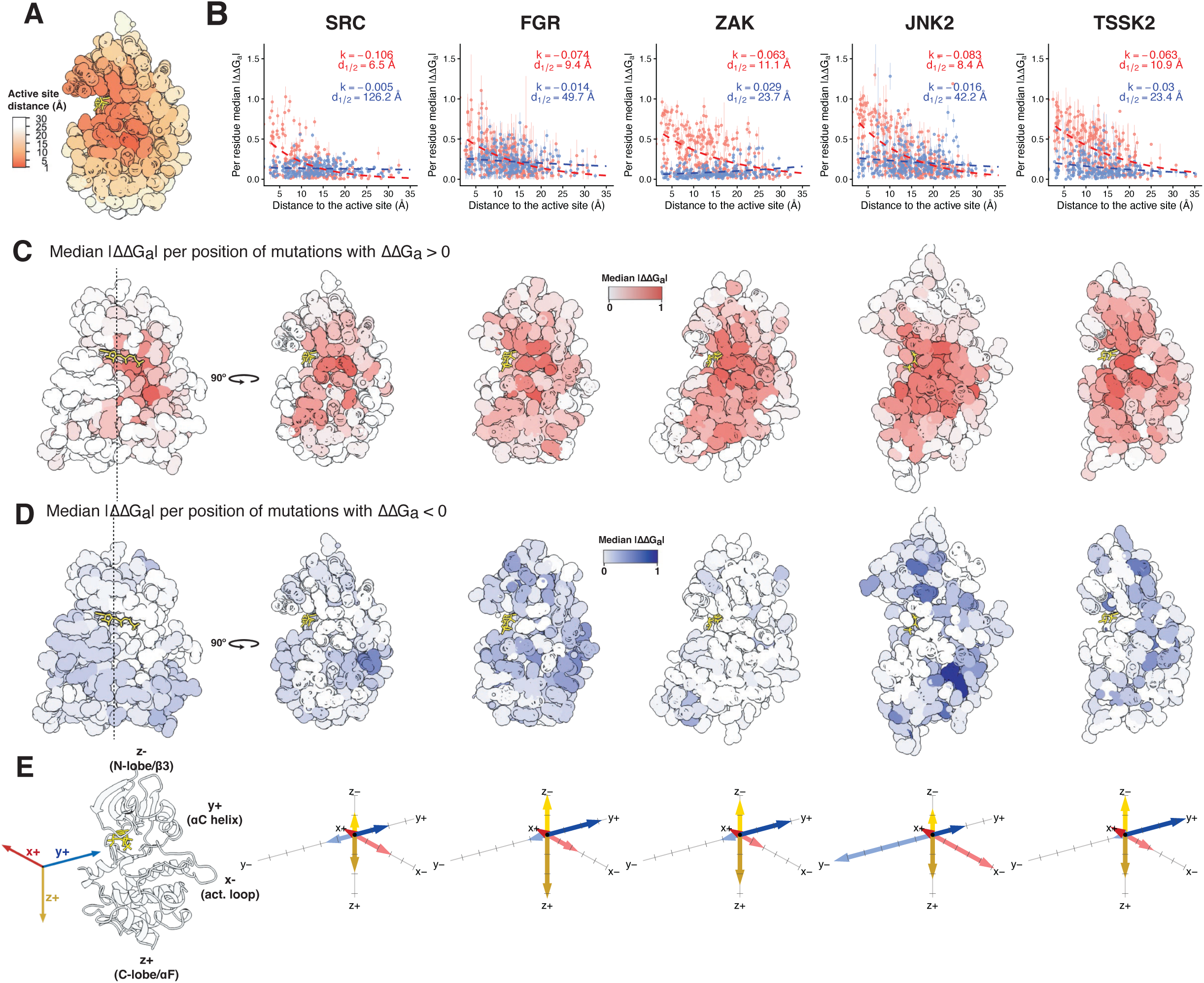
Distance dependence and anisotropy of allosteric effects on activity. (A) SRC structure with each position coloured by its distance to the active site (Å). (B) Per-residue median |ΔΔGa| versus the minimum distance to the active site (ATP and catalytic aspartate) in each kinase, for inhibitory (ΔΔGa > 0, red) and activating (ΔΔGa < 0, blue) mutations. Error bars, standard error of the median. Dashed curves show exponential decay fits (y = a · e^kx^); d½ = ln(2)/|k|. (C) Per-position median |ΔΔGa| of inhibitory mutations (ΔΔGa > 0) mapped onto the structure of each kinase. SRC is shown in two views related by a 90° rotation. (D) As in (C) for activating mutations (ΔΔGa < 0). (E) Directional anisotropy of the activity decay. Left, the x/y/z coordinate frame on the SRC structure (y+, αC helix; x-, activation loop; z+, C-lobe and αF; z-, N-lobe and β3). Right, for each kinase, arrows along the six directions (x±, y±, z±) with length proportional to the allosteric half-distance d½ in that direction. The axis tick marks are spaced every 10 Å. ATP is shown as yellow sticks in all panels.

### Anisotropic allostery

For each kinase, distance-dependent allostery explains a substantial amount of the variance in ΔΔGa for inactivating mutations, but only a minority - on average 29% (range: 23% for ZAK to 36% for JNK2). At any particular distance from the active site there is substantial variation in both the average allosteric effect of mutations at each residue (Figure S6A, top) and the effects of individual mutations at each site (Figure S6A, bottom). To test if allosteric decay differs in different spatial directions, we quantified allosteric decay in three orthogonal directions from the active site (x, y, z, defined in the SRC structure [PDB = 2SRC], Figure 4E). In all five proteins, allosteric coupling is strongest towards helix αC (y+ direction, median d½ = 26.1 Å). Coupling is intermediate along the z-axis in both directions (z+ direction towards the C-lobe, median d½ = 18.6 Å); z-direction towards the N-lobe, median d½ = 14.1 Å). Coupling is weaker along the x-axis towards helix αD (and the SH2/SH3 regulatory-domain interface in the tyrosine kinases FGR and SRC) (x+ direction, median d½ = 4.8 Å).

However, the strength of allostery in a given direction also varies across the kinases (Figure 4E). Towards the αF/C-lobe (z+), FGR has stronger coupling than SRC (d½ = 28.6 Å, 95% CI 23.8 to 35.7, versus 15.2 Å, 95% CI 13.8 to 16.9; z-test on decay constants, z = 6.3, p < 0.05). In the αC-helix direction (y+), allostery is stronger in ZAK than in JNK2 (d½ = 26.1 Å, 95% CI 20.5 to 36.1, versus 15.1 Å, 95% CI 13.0 to 18.1; z = 3.6, p < 0.05), while towards the activation loop (x-), JNK2 has the strongest coupling (d½ = 46.4 Å, 95% CI 33.3 to 76.4) and ZAK the weakest (d½ = 14.9 Å, 95% CI 12.2 to 19.0; JNK2 versus ZAK, z = 5.3, p < 0.05). The strong activation-loop coupling in JNK2 is consistent with the control of MAPK activity by activation-loop phosphorylation^54^.

Allosteric decay rates do not differ between β-strands and α-helices in any kinase (β-strand d½ = 7.5, 12.0 and 5.9 Å versus α-helix d½ = 11.8, 22.2 and 9.2 Å in SRC, FGR and JNK2; z-test on decay constants, p > 0.05) (Figure S6B). Particular types of mutation are more likely to be allosteric, with inhibitory allosteric mutations enriched at isoleucine, tyrosine, histidine and valine wild-type residues (OR = 2.05, 1.95, 1.89 and 1.50; all FDR < 0.1) and for substitutions to proline, arginine, glycine and tryptophan (OR = 1.98, 1.80, 1.37 and 1.33; all FDR < 0.1) (Figures S6E and S6H).

### Activating mutations are structurally dispersed and kinase-specific

We next considered activating mutations occurring outside the kinase active sites. In contrast to inhibitory mutations, activating mutations are not enriched near the active site but are dispersed throughout the kinase domain (Figures 4B and 4D). They are, however, enriched at particular residues (one-sided FET, FDR < 0.1) (Figures 5A, 5C and S7A). The number of activating mutation-enriched sites per kinase ranges between 2 (TSSK2) and 23 (JNK2), with a median of 9. Activating sites are, however, different for each kinase, with 43 of the 47 residues enriched only in a single protein (Figures 5C and S7A). Three of the four shared residues are enriched in both SRC and FGR, the two most closely related kinases (alignment positions 141 and 147 in the αE helix and position 220 in the αF helix). In addition, position 221 in the αF helix is enriched in both SRC and JNK2. A further 7 of the 47 residues are enriched for activating mutations in one kinase and for inactivating mutations in another (positions 79, 150, 156, 176, 247, 260 and 329), further highlighting the divergence in mutational effects. In total, 860 of the 946 distinct activating mutations are only activating in a single kinase and none activate in all five (Figure S6G).

**Figure 5.**
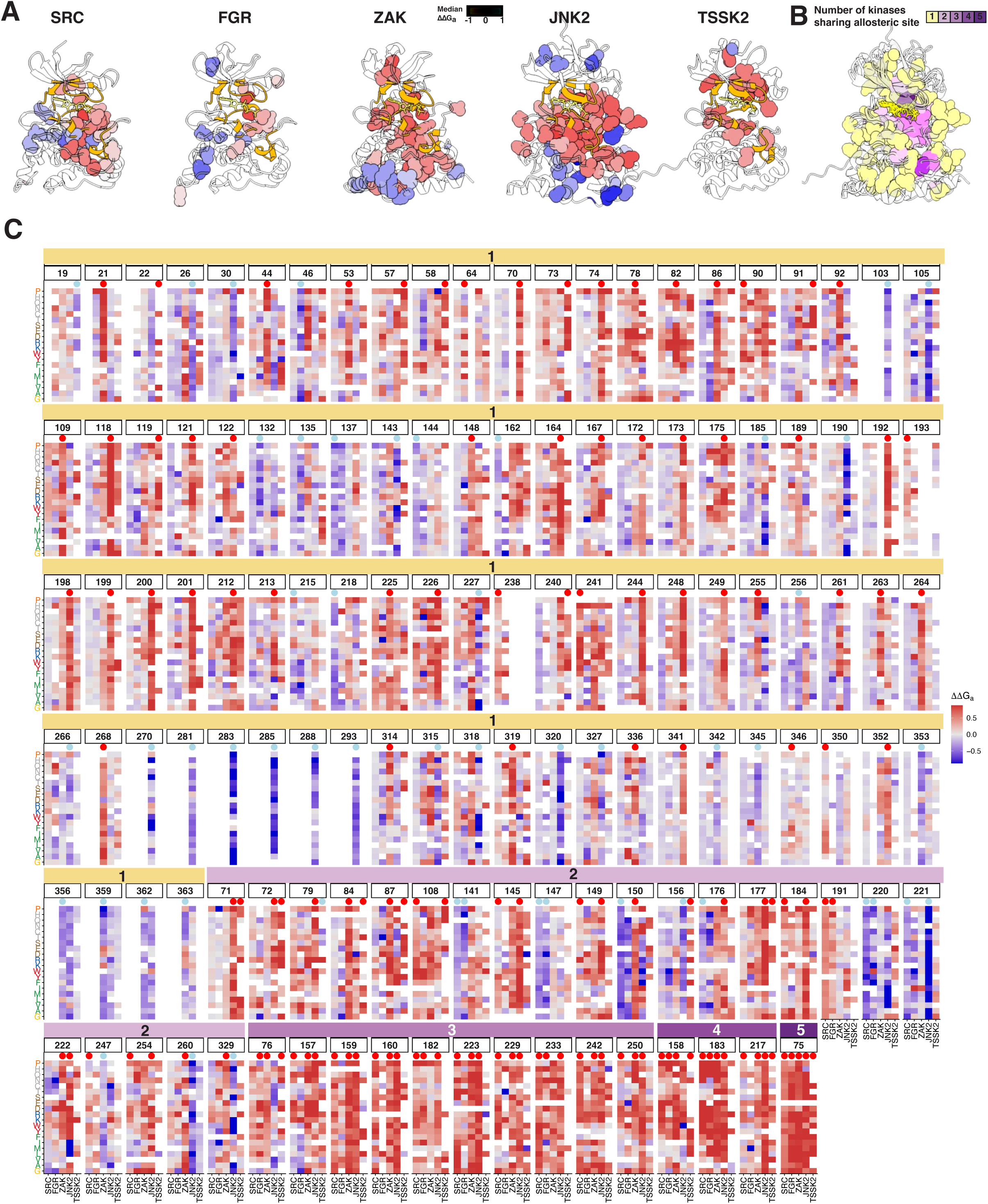
**Major allosteric sites across five kinases**. (A) Major allosteric sites on each kinase structure, shown as spheres coloured by per-residue median ΔΔGa. ATP is shown as yellow sticks. (B) The five kinase structures superimposed, with major allosteric positions coloured by the number of kinases in which the aligned position is an allosteric site (1 to 5, yellow to purple). (C) For each major allosteric site, ΔΔGa of all substitutions (rows, mutant amino acid) across the five kinases (columns), grouped by the number of kinases sharing the hotspot (1 to 5). Red and blue dots mark positions enriched for inhibitory and activating allosteric mutations, respectively.

In SRC, JNK2, and ZAK, activating mutations are more frequent in the C-lobe (OR = 3.9, 1.3, and 18, respectively, all p < 0.05), but in TSSK2 activating mutations are more frequent in the N-lobe (OR = 17, p < 0.05) (Figure S6C). In SRC and FGR activating mutations are most enriched in the αE helix (OR = 4.1 and 2.2, respectively; FDR < 0.1), whereas in JNK2 they are most enriched in the αI helix (OR = 2.6, FDR < 0.1), in ZAK in the αG helix (OR = 14, FDR < 0.1), and in TSSK2 in the β1 strand (OR = 14, FDR < 0.1) (Figure S6F). Considering all five kinases, mutations to tryptophan, phenylalanine and glycine are enriched for activating mutations (OR = 1.61, 1.55, 1.44 respectively, FDR<0.1). Activating mutations are more likely to occur at leucine residues across all kinases (OR = 1.29, FDR = 0.07). However, some wild type residues are enriched in activating mutations only in individual kinases, for example mutations at alanine in SRC (OR = 2.4), phenylalanine in JNK2 (OR = 2.28), and leucine in ZAK (OR = 2.56) (FET, all FDR < 0.1) (Figures S6E and S6H).

Activating mutations are thus rare, structurally dispersed, and idiosyncratic to each kinase.

### Major allosteric sites

Considering both inactivating and activating mutations, 185 residues at 129 positions are enriched for allosteric mutations in at least one kinase, residues we refer to as ‘major allosteric sites’ (Figures 5A, 5C and S7A). Most major allosteric sites are enriched for inactivating mutations (134 of 185 residues; 89 different sites). 51 residues at 47 different sites are enriched for activating residues. The major allosteric sites form a gradient from conserved to kinase-specific. A single residue - the conserved αC-helix glutamate - is a major allosteric site in all five kinases. Three residues are major allosteric sites in four kinases (158, 183, 217), 10 in three, and 23 in two. 92 sites are enriched for allosteric mutations in a single kinase (Figure 5C). Of the 93 residues at shared major allosteric sites (enriched in two or more kinases), 81 (87%) are located in the core and 12 (13%) on the surface. In contrast, of the 92 kinase-specific major allosteric sites, 26 (28%) are located on the surface (OR = 2.6, FET p < 0.05) (Figures 5B and S7B). Indeed, considering all non-active site positions, mutational effects on activity are better correlated in the core compared to the surface (Wilcoxon p < 0.05, median Spearman ρ= 0.081 for core sites, and median ρ = 0.015 for surface sites) (Figure S7C).

In all five kinases, the major allosteric sites are spatially clustered (p < 0.1, permutation test for within-8 Å contact-density) (Figure 5A). However they are in different locations in each protein. In JNK2 the major allosteric sites are concentrated in the activation segment (activation and catalytic loops) and in the C-lobe substrate-binding helices (αF, αC and αG helices). In ZAK the major allosteric sites span both lobes, including in the αC helix, the β4 strand and the C-lobe αF/αG helices. In TSSK2 they concentrate in the N-lobe, around the αC helix and the β3-β5 sheet. In SRC the major allosteric sites cluster in the C-lobe αE and αF helices and the catalytic/activation loops, and in FGR they are concentrated in the αE helix and activation loop. These sites are predominantly inactivating. The activating sites are fewer (2 to 23 per kinase) and, where testable (≥5 sites), cluster within a subset of these regions: in SRC on the αE and αF surface bound by the SH2 domain (11 sites, p < 0.1), in JNK2 on the αH helix and activation loop (23 sites, p < 0.1), and in ZAK on the αG helix (9 sites, p < 0.1). They do not cluster in FGR (6 sites, p = 0.21), and TSSK2 has too few (2) to test.

Within each kinase there is therefore a gradient of more conserved major allosteric sites that are closer to the active site and more protein-specific major allosteric sites towards specific regions of the protein surface (Figure 5A and 5B).

### Conservation and divergence of allosteric surfaces

For physiological regulation and the development of therapeutics, the solvent-accessible surface of a protein is particularly important. We therefore further compared the effects of mutations across the surfaces of the five kinase domains (Supplementary Video 1-5). Across the ‘front’, ‘back’, ‘left’ and ‘right’ surfaces of the two most related kinases, SRC and FGR, the median effects of mutations are quite similar (Figure 6) (r = 0.75 for median ΔΔGa across all surface sites). In each, activating mutations cluster on the C-lobe below the ATP cleft, over the αE and αF helices and the β4 strand (right and back faces), with a shared activating allosteric site on αF (aligned position 220, ΔΔGa = -0.56 kcal/mol in SRC and -0.32 kcal/mol in FGR). Inhibitory mutations are more sparse, located mainly in the activation loop and the N-lobe β-sheet (β1 and β2 strands).

**Figure 6.**
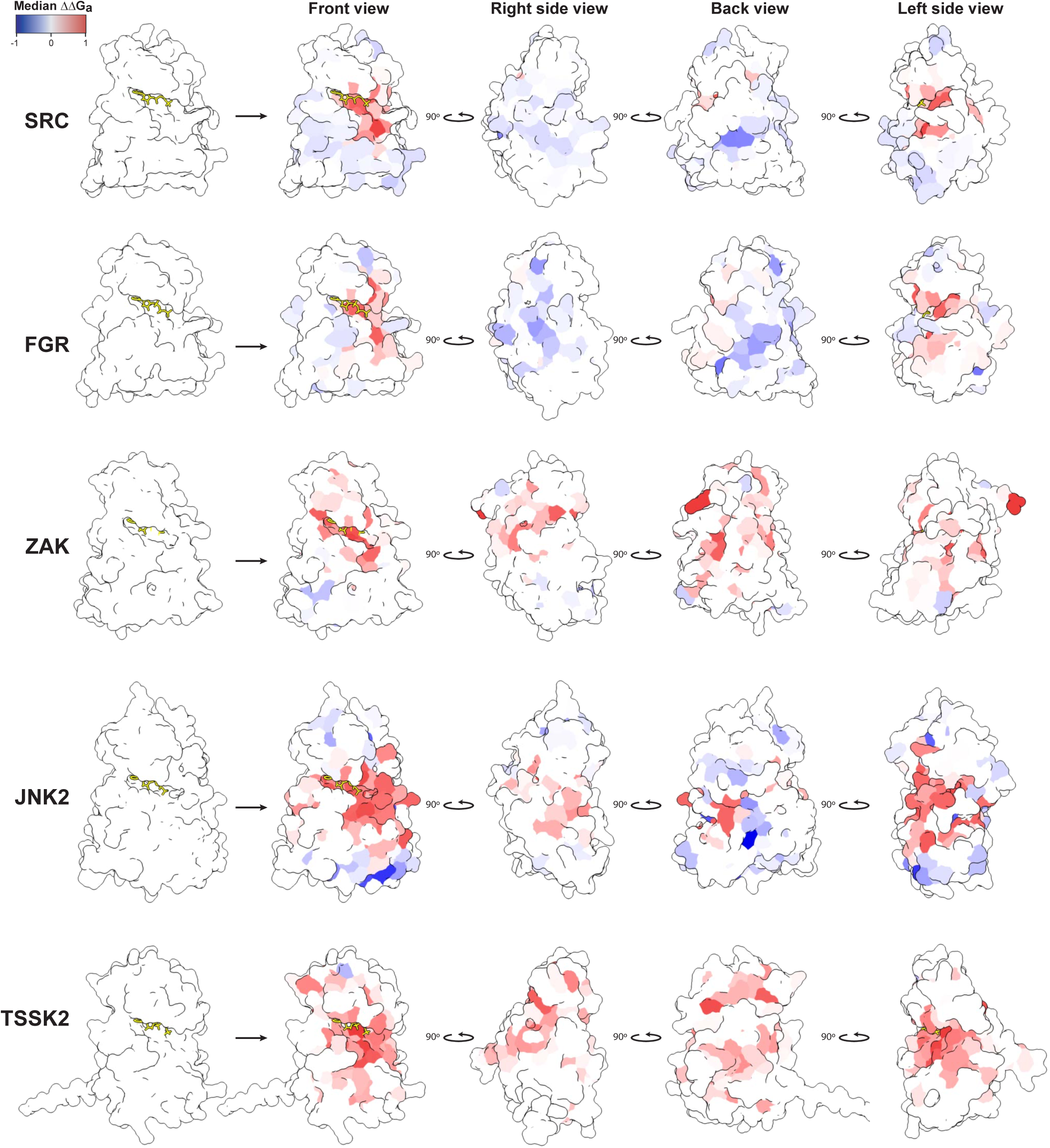
Comparative allosteric surfaces. Each row is one kinase (SRC, FGR, ZAK, JNK2 and TSSK2). The first column shows the kinase domain with the ATP-site ligand (yellow sticks), uncoloured and in the same orientation as the front view, for reference. The next four columns show the surface coloured by the per-position median ΔΔGa, rotated by 90° about the vertical axis at each step: front, right side, back and left side views.

Comparing more distant kinases however, reveals substantial divergence in ΔΔGa across the surface (r = 0.66, 0.52 and 0.40, comparing SRC to ZAK, TSSK2 and JNK2, respectively). ZAK and TSSK2 are inhibition-biased, with few positions enriched for activating mutations (1 and 4, versus 30 and 19 enriched for inactivating), but their inhibitory surfaces are organised differently (Figure 6): in ZAK they concentrate on the right and back faces, running across the activation loop (aligned 179 to 202, ΔΔGa up to 0.90 kcal/mol), the catalytic loop (Arg160) and the β4 strand, whereas in TSSK2 they are spread more evenly across all four faces. The JNK2 surface has residues enriched for mutations with both signs, with near-neutral median effects on all four faces (median ΔΔGa = -0.04 to 0.00 kcal/mol): it shares the SRC and FGR activating allosteric site on αF (aligned 221, ΔΔGa = -1.28 kcal/mol), but its inhibitory allosteric sites concentrate on the activation loop (a cluster at aligned 184, 192, 198, 200 and 201; ΔΔGa = 0.74 to 0.82 kcal/mol) including the activation-loop phosphosite, and on the αF and αG helices (Tyr217 ΔΔGa = 0.82 kcal/mol; Ile248 ΔΔGa = 0.87 kcal/mol). The allosteric surfaces are thus quite diverged, with mutations more frequently activating in SRC and FGR, more frequently inhibitory in ZAK and TSSK2, and more balanced in JNK2.

### A cluster of allosteric pockets conserved in all five kinases

The high structural conservation of the ATP-binding pocket across kinases makes the development of kinase-specific orthosteric inhibitors very challenging^65^. Targeting allosteric sites can enable the development of more specific inhibitors, provided these sites have lower structural or functional conservation^66^. Targeting allosteric sites can also overcome drug resistance mutations^17,18^ and allows the development of kinase activators^23,67^. For most kinases, however, we do not know which potentially-druggable surface pockets are allosteric and which are not. We also do not know how conserved or divergent this coupling is in different members of the protein family. We therefore used our comprehensive allosteric maps to identify and compare the functional pockets in each of the five kinases.

We used FTmap^68^ to identify a total of 37 distinct fragment-sized pockets across the five proteins (Figures 7A, 7C and S8A, S9 and Table S4). 16 of these pockets correspond to 10 named pockets in the Kinase Atlas database: the ATP, DFG, MT3, MPP, PIF, DRS, PDIG, DEF, LBP, and EDI pockets^69^. For each pocket in each kinase we quantified its negative and positive allostery as the enrichment of inhibitory and activating allosteric mutations, respectively (Figures 7A and S8C).

**Figure 7.**
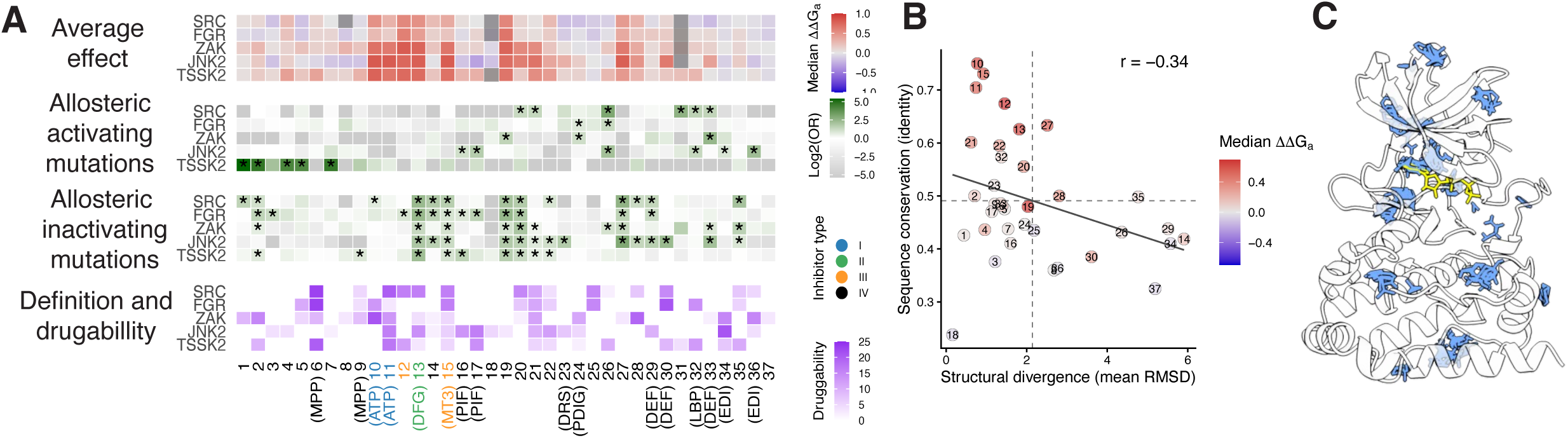
Allosteric pockets across five kinases. (A) Summary of the 37 global pockets (columns, with classic pocket names in parentheses) in each kinase (rows). Top panel, median ΔΔGa of the pocket residues; grey, pocket with < 4 residues in that kinase. Second and third panels, enrichment (log2 odds ratio) of activating and inactivating allosteric mutations in each pocket relative to the whole kinase domain (one-sided hypergeometric test; asterisks if FDR < 0.1). Bottom panel, FTMap druggability score of the pocket in each kinase. Pocket-name labels are coloured by inhibitor type. (B) Sequence conservation (mean residue identity across the five kinases) against structural divergence (mean pairwise Cα RMSD) for each pocket. Points are labelled by pocket number and coloured by ΔΔGa (per-kinase pocket median, averaged across kinases). Dashed lines mark the means and the solid line marks the linear fit. (C) FTMap probe clusters (blue) on the ZAK structure. ATP is shown as yellow sticks.

Strikingly, 30 out of 37 pockets are enriched for allosteric mutations in at least one kinase, reflecting the widespread allosteric potential of kinase surfaces (Figure 7A). Five pockets are enriched for inhibitory mutations in all five kinases: the ATP binding site (pockets 10 and 11), the DFG pocket (pocket 13, Figure 8), where Type-II inhibitors bind^70^, and three additional pockets (Figure 7A). These functionally conserved inhibitory pockets are the MT3 pocket (15), where Type-III inhibitors including trametinib and cobimetinib bind in MEK1 and MEK2 and disrupt the β3-αC salt bridge^71^, and the uncatalogued pockets 19 and 20. All three pockets are located in the cleft between the N and C lobes and all are adjacent to residues that contact ATP and directly participate in catalysis. The DFG pocket, MT3 pocket and pockets 19 and 20 have lower structural and sequence conservation than the orthosteric site (Figures 7B and S8B) (Cα RMSD 1.82, 0.92, 2.03 and 1.93 Å respectively, versus 0.74 and 0.77 Å for the two ATP pockets), but are more structurally conserved than the average across all pockets (mean RMSD 2.14 Å across all 36 pockets; Figure 7B). The enrichment of inhibitory mutations in these pockets in all five kinases is striking and suggests they may constitute a functionally-conserved inhibitory surface to target in many - and perhaps all - protein kinases.

**Figure 8.**
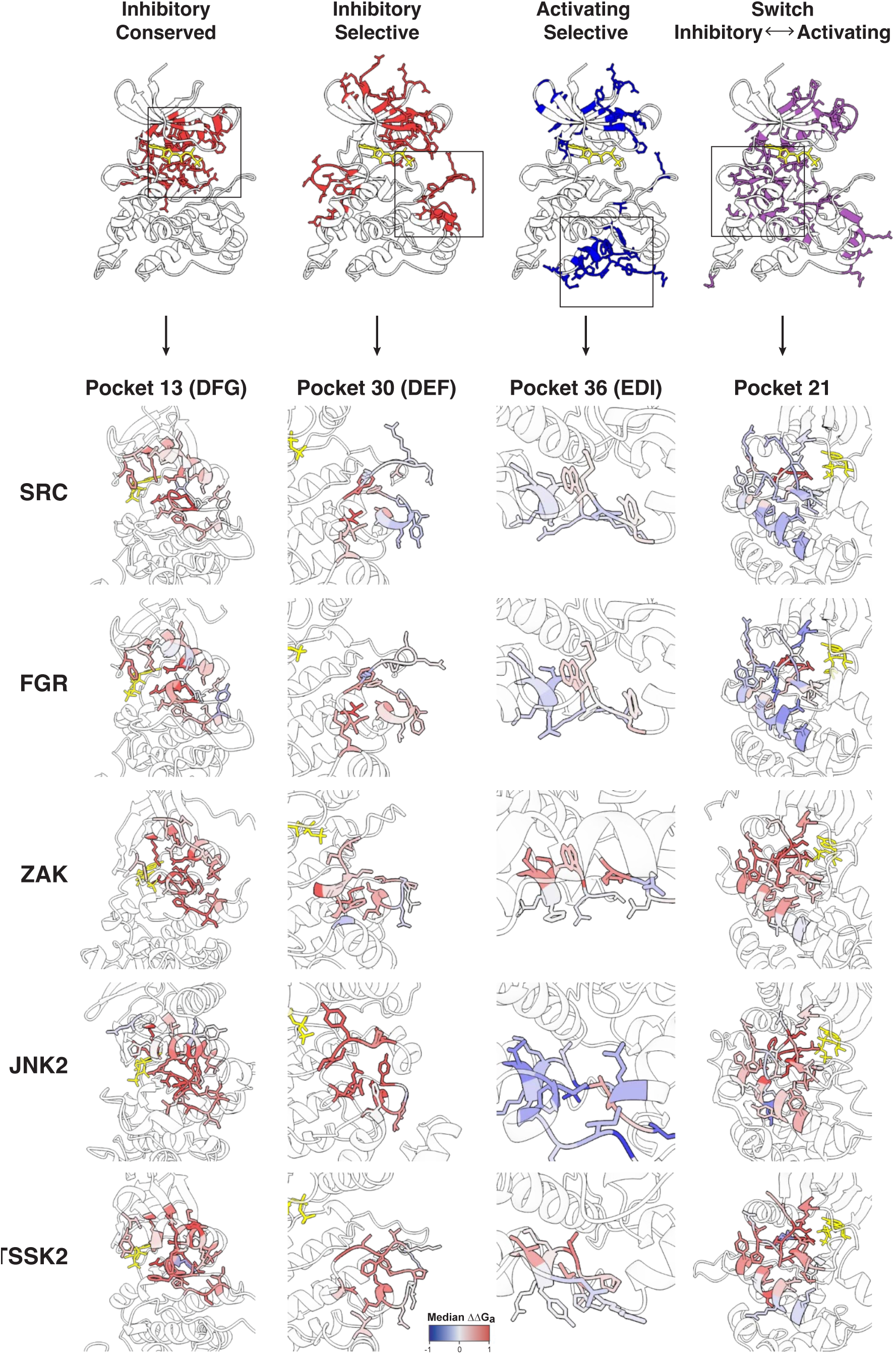
Conserved, kinase-specific and switching allosteric pockets. Top, the four pocket categories mapped onto the SRC structure: inhibitory pockets conserved in all five kinases, inhibitory pockets specific to one kinase, activating pockets specific to one kinase, and switching pockets. Boxes mark the example pocket shown below each category. Bottom, close-up of one example pocket per category in each kinase: Pocket residues are coloured by median ΔΔGa. ATP is shown as yellow sticks.

### Each kinase has a unique set of inhibitory pockets to target

Beyond the ATP pockets and the four conserved inhibitory pockets, each kinase has a different repertoire of allosteric pockets. The total number of additional pockets enriched for allosteric mutations ranges between 7 (ZAK) and 16 (JNK2), with 5 to 10 inhibitory and 2 to 6 activating pockets per kinase (Figure 7A). Two pockets are enriched for inhibitory mutations in four kinases: pocket 2, which is located between the αC helix and the β4 strand of the N-lobe, in SRC, FGR, ZAK and TSSK2 (OR 1.5-2.7, FDR < 0.1) and pocket 27, which spans the catalytic and activation loops of the C-lobe, in SRC, FGR, ZAK and JNK2 (OR 2.5-7.6, FDR < 0.1). A further five pockets are enriched for inactivating mutations in three kinases each: pocket 14 (central activation loop, at the phosphosite and P+1 loop), pocket 21 (N-lobe β4 strand and αC helix, the αE helix, and the DFG motif starting the activation loop), pocket 22 (catalytic loop and αD helix, reaching the P+1 loop), the DEF pocket (pocket 29; the phosphosite and APE end of the activation loop, with the αG helix) and pocket 35 (αF, αG and αH helices) (OR 1.3-4.1, FDR < 0.1).

We next identified high-confidence kinase-specific inhibitory pockets as those enriched in inhibitory mutations in a single kinase (FET FDR<0.1), and whose distribution of ΔΔGa is shifted towards inhibition in all pairwise comparison to the homologous pockets in other kinases (baseline-corrected Wilcoxon rank sum test FDR<0.1, see Methods) (Figure S8D). We identified three high-confidence kinase-specific pockets, two in JNK2 and one in SRC. The two JNK2-specific pockets are both known MAP kinase substrate-recruitment surfaces: the D-motif docking site (DRS; pocket 23) and the FXF docking site (DEF; pocket 30; Figure 8), enriched for inhibitory mutations (OR = 3.3 and 5)^58,72,73^. The SRC-specific pocket is uncatalogued pocket 1 (SRC, OR = 2.7), an N-lobe site at the αC helix and β4 strand adjacent to pocket 2.

The energetic maps thus reveal that all five of the kinases have multiple potentially druggable functional pockets to target, including pockets functionally active in all five proteins and functional pockets specific to single kinases only.

### Selectively targeting highly-related kinases

A frequent challenge in drug development is modulating one protein without affecting a highly-related homolog. FGR and SRC are two closely related kinases and they indeed have the two most similar functional landscapes in our data, with highly correlated energy landscapes (ΔΔGa r = 0.79, compared to median ΔΔGa r = 0.49 across the five proteins. In our maps FGR and SRC share eight inhibitory allosteric pockets (2, 13, 14, 15, 19, 20, 27 and 29), including four that are conserved across the five kinases (the DFG pocket 13, MT3 pocket 15, and pockets 19 and 20). However, pockets 1, 22, 28, and 35 are enriched for inactivating allosteric mutations in SRC (OR = 2.7, 1.9, 2.3, and 4.1, respectively; all FDR < 0.1) but not in FGR (OR = 0.9, 0.8, 1.0, and 1.2, all FDR > 0.1). These functional pockets are therefore candidate sites for the development of more specific inhibitors, potentially allowing the selective targeting of SRC over FGR.

### Activation pockets

In total, 17 pockets are enriched for activating mutations (FET, FDR < 0.1). However, 13 of these pockets are only enriched for activating mutations in a single kinase, making them much more kinase-specific than the inhibitory pockets (only 9 of 22 are kinase-specific). Pocket 26, an uncatalogued C-lobe pocket on the αE/αF face, is enriched for activating mutations in three kinases (Figure 8): SRC, FGR and JNK2 (OR 2.4-8.1, FDR < 0.1), suggesting a site positively coupled to the active site across both tyrosine kinases (SRC, FGR) and MAP kinases (JNK2). Three further pockets are enriched for activating mutations in two kinases each. Pocket 32 is homologous to the LBP (lipid-binding) pocket, formed by the MAP kinase insert in the C-lobe of p38α^75^ and it it is enriched in SRC and JNK2 (OR 2.5-2.9, FDR < 0.1). Pocket 33 (DEF) is enriched in SRC and ZAK (OR 2.7-5.4, FDR < 0.1). Pocket 24 is a C-lobe pocket homologous to the PDIG substrate-recognition site, which is targeted by allosteric inhibitors in Chk1^76,77^ and also found in PIM1, DAPK and the JNK kinases^69^), and it is enriched in FGR and ZAK (OR 1.7-2.2) (FDR < 0.1).

We identified three high-confidence kinase-specific activating pockets as those enriched in activating mutations in a single kinase (FET FDR<0.1), and whose ΔΔGa distributions are shifted towards activating mutations in pairwise comparisons (baseline-corrected Wilcoxon rank sum test FDR<0.1) (Figure S8D). These are pocket 1 in TSSK2 (OR = 41.9, FDR < 0.1), located between the PIF and MPP sites on the αC/β4 face, and pockets 17 and 36 in JNK2. Pocket 17 (OR = 3.3, FDR < 0.1) is the PIF site, a regulatory hydrophobic motif docking groove on the N-lobe. Pocket 36 (OR = 4.7, FDR < 0.1) is the C-lobe EDI site, the EGFR dimerization interface, that in EGFR drives allosteric activation by asymmetric dimerization^78^.

### Pockets that switch from activation to inhibition

Interestingly, mutations in several pockets have opposite effects in different kinases, with seven pockets enriched for activating mutations in one kinase but inactivating mutations in another. Four of these lie on the αC helix and the β3-β5 strands of the N-lobe. For example, mutations in the PIF pocket (pocket 16) are activating in JNK2 (OR 2.1 and 3.3, both FDR <0.1) but inhibitory in TSSK2 (OR = 2.8, FDR <0.1). This diversity in the outcome of mutations in the PIF pocket mirrors that of allosteric modulators targeting this pocket, which have been shown to elicit both activating and inactivating effects^79^.

Mutations in the uncatalogued αC/β4 pocket 2 are activating in TSSK2 (OR = 26.9, FDR <0.1) but inhibitory in SRC, FGR and ZAK (OR = 1.5-2.7, all FDR <0.1). The other three ‘switch’ pockets sit at distinct positions. Mutations in pocket 21 (β4 strand, αE helix and activation loop, Figure 8) are activating in SRC (OR = 2.6, FDR <0.1) but inhibitory in ZAK, JNK2 and TSSK2 (OR = 1.3-1.6, all FDR <0.1); mutations in pocket 26 (αE and αF helices of the C-lobe) are activating in SRC, FGR and JNK2 (OR = 2.4-8.1, all FDR <0.1) but inhibitory in ZAK (OR = 2.0, FDR <0.1); and mutations in the DEF pocket (pocket 33; αG helix and activation loop) are activating in SRC and ZAK (OR = 2.7 and 5.4, both FDR <0.1) but inhibitory in JNK2 (OR = 4.3, FDR < 0.1).

Finally, two pockets are enriched for both activating and inhibitory mutations in the same kinase: pocket 2 (αC helix and β4 strand of the N-lobe) in TSSK2 (activating OR=26.9, inhibitory OR = 1.7) and the DEF pocket (pocket 33) in ZAK (activating OR = 5.3, inhibitory OR=1.8). In TSSK2, pocket 2 contains an activating position (aligned 19, median ΔΔGa = -0.37 kcal/mol) among several inhibitory residues (aligned 91, 82, 76, 93, 92, 18 and 89; median ΔΔGa = 0.22 to 0.85 kcal/mol), and similarly in ZAK the DEF pocket pairs an activating position (aligned 256, median ΔΔGa = -0.29 kcal/mol) with several inhibitory residues (aligned 255, 212 and 248; median ΔΔGa = 0.21 to 0.48 kcal/mol). The functional effect of perturbations in these pockets thus depends on which residue is perturbed.

In summary, our data show that each kinase has a distinct set of functional pockets to potentially therapeutically target. These include inhibitory pockets close to the orthosteric site that are functionally-active in all five enzymes, as well as more distant pockets where function is more kinase-specific. In addition, all five kinases have at least one distal pocket where mutations activate the enzyme, identifying potential sites for the development of kinase activators (Figure 8). For seven activation pockets, mutations are actually inhibitory in a different kinase. The functional pocket map of each kinase is therefore unique, with a distinct allosteric surface to regulate and therapeutically target. This divergence highlights the importance of constructing a protein-specific allosteric map for each kinase.

## Discussion

We have presented here complete energetic and allosteric maps for five protein kinases and a comparative analysis of allostery in an enzyme family. By measuring the activity and abundance of >160,000 kinase domain variants using the KINASE-MAPS platform we quantified the effects of all mutations in all positions on both fold stability and the enzymatic activity of five different kinases. These complete energetic and allosteric maps provide a number of important insights into the functional landscapes of protein kinases and how to potentially modulate them therapeutically.

In all five proteins, inhibitory mutations show a conserved distance-dependent decay away from the active site. The allosteric architecture is not symmetric though, with mutations in particular regions more likely to inhibit activity and this allosteric anisotropy is different across the five proteins. For activating mutations the architecture is very different, with no enrichment close to the active site and each kinase having a distinct set of activation sites. In all five kinase domains the network of allosterically active residues extends to the surface, and all five proteins have multiple allosterically active surface pockets to potentially therapeutically target.

For kinase inhibition, a functionally-conserved cluster of inhibitory pockets exists between the N- and C-lobe. It is likely that these inhibitory interlobe pockets are allosteric in most — and perhaps all — kinases, with the design of kinase-selective allosteric inhibitors requiring the exploitation of structural divergence^16^. In addition, however, all five kinases have additional inhibitory pockets to target further away from the active site, with these more distal pockets less functionally conserved across the five proteins. Even the highly similar kinases FGR and SRC have differences in functional pockets, potentially facilitating the development of more selective inhibitors^33^.

All five kinases also have pockets where mutations consistently activate the protein, highlighting the potential for the development of kinase activators. These activation pockets are less functionally conserved than the inhibitory pockets; kinase activators are therefore more likely to be protein-specific. Interestingly, in two pockets across the five proteins, different mutations activate and inhibit kinase activity, illustrating the importance of the exact perturbation in a pocket. Finally, multiple pockets are enriched for activating mutations in one kinase but for inhibitory mutations in another, further illustrating how the functional landscape of a conserved protein structural fold qualitatively changes during evolution.

Our energy landscapes also reveal alterations in the importance of residues for fold stability across the kinase domain fold. The conservation of mutational effects on fold stability has been little investigated in larger protein domains and it will be interesting in future work to explore this more generally and to understand mechanistically why the energetic effects of mutations diverge during evolution.

Most importantly, however, our data show that proteins with the same 3D-fold do not have the same energetic or allosteric structure. Each of the five kinases has an extensive allosteric network that shares certain principles, but the allosteric structure and surface is also unique to that protein. The large allosteric network of each protein should empower the design and discovery of kinase modulators and the divergence of the allosteric structure should allow the development of protein-specific inhibitors and activators.

It is likely that architecture that we describe here — a conserved catalytic core overlaid with divergent regulatory control — is a general principle in the evolution of allosteric regulation in protein kinases^80^, and across other protein families^59^. We believe it is therefore critical to build an experimental energetic and allosteric map for each human protein of therapeutic interest. Mapping the ‘functional pocketomes’ of kinases and other proteins will help unlock the identification of functional and selective pockets to therapeutically target. Only with protein-specific experimental maps of allostery and functional pockets will we be able to identify the best sites to target to selectively modulate each protein.

## Methods

### Media

- LB: Bacto-tryptone (10 g/L), yeast extract (5 g/L), and NaCl (10 g/L). Autoclaved for 20 min at 120°C
- YPD: glucose (20 g/L), peptone (20 g/L), yeast extract (10 g/L). Autoclaved for 20 min at 120°C
- YPDA: glucose (20 g/L), peptone (20 g/L), yeast extract (10 g/L), and adenine sulphate (40 mg/L). Autoclaved for 20 min at 120°C
- SORB: 1 M sorbitol, 100 mM LiOAc, 10 mM Tris-HCl (pH 8.0), and 1 mM EDTA.
- Recovery medium: YPD plus 0.5 M sorbitol. Filter sterilized.
- Plate mixture: 40% PEG 3350, 100 mM LiOAc, 10 mM Tris-HCl (pH 8.0), and 1 mM EDTA (pH 8.0).Filter sterilized.
- SC-URA: yeast nitrogen base without amino acids (6.7 g/L), glucose (20 g/L), and complete supplement mixture drop-out without uracil (0.77 g/L). Filter sterilised.
- SC-URA + 2% raffinose + 0.1% glucose: yeast nitrogen base without amino acids (6.7 g/L), raffinose (20 g/L), glucose (1 g/L), and complete supplement mixture drop-out without uracil (0.77 g/L). Filter sterilised.
- SC-URA + 2% galactose + 0.1% glucose: yeast nitrogen base without amino acids (6.7 g/L), galactose (20 g/L), glucose (1 g/L), and complete supplement mixture drop-out without uracil (0.77 g/L). Filter sterilised.
- SC-URA-ADE: yeast nitrogen base without amino acids (6.7 g/L), glucose (20 g/L), complete supplement mixture drop-out without uracil and adenine (0.76 g/L). Filter sterilised.
- MTX competition medium: SC-URA-ADE plus methotrexate (200 µg/mL; BioShop Canada Inc., Canada) and 2% DMSO.
- DNA extraction buffer: 2% Triton X-100, 1% SDS, 100 mM NaCl, 10 mM Tris-HCl (pH 8.0), and 1 mM EDTA (pH 8.0).

### Activity-dependent toxicity assay

We selected four kinases reported as toxic in yeast by Kim et al. 2020^81^. We amplified them from the ORFeome collection and cloned each one separately, by Gibson Assembly (NEB), into the plasmid used to evaluate toxicity (pCF85). We ordered an oPool from Integrated DNA Technologies (IDT) containing all single mutations of the most N-terminal region of each kinase domain, encoded with NNK codons and corresponding to 59 to 65 positions depending on the kinase, based on their multiple sequence alignment (Table S3). We amplified each kinase oPool separately by PCR and cloned it, by Gibson Assembly (NEB), into the corresponding toxicity-evaluation plasmid, previously linearised.

Each of the four libraries constructed in *Escherichia coli* (*E. coli*) was transformed into yeast (BY4741 strain) in triplicate, at a coverage of at least 100 clones per variant. These were subjected to an activity-dependent toxicity selection, in which each kinase was overexpressed from a strong galactose-inducible promoter (toxicity selection). After selection, cells were harvested and plasmid DNA was extracted. To prepare the four NNK libraries for Illumina sequencing, two rounds of PCR amplification were performed to add the Illumina adapter sequences and a unique index to each sample. The resulting amplicons were sequenced by the CRG Genomics Core Facility using NextSeq 2000 (Illumina) paired-end 2×150 bp sequencing. Sequencing data were processed with DiMSum^82^.

### Variant library design and construction

To cover the full kinase domain of the four kinases, we mutagenised them using SUNi mutagenesis^44^. Each full-length kinase from the ORFeome was cloned into a generic mutagenesis plasmid (pCF121) using the NheI and NotI restriction enzymes (NEB).

To obtain double mutants, we first cloned, by NEBuilder HiFi DNA Assembly (NEB), oPools from IDT encoding WWC at two specific codon positions, giving a total of 8 nucleotide variants per kinase as starting templates for the mutagenesis. Oligonucleotides designed to encode all amino acid substitutions (using NNK or NNS degenerate codons) were also ordered as an oPool from IDT. The standard SUNi mutagenesis protocol was scaled up fivefold to obtain sufficient transformant coverage per variant. A total of 4.5 μl of the final product was electroporated into three cuvettes, each containing 25 μl of NEB 10β High-efficiency Electrocompetent *E. coli* cells, followed by 1 h recovery in 2 ml of super optimal broth with catabolite repression (SOC) medium. 0.1% of the recovery product was plated onto LB agar with spectinomycin, and the rest was inoculated into 100 ml of LB liquid with spectinomycin. For each kinase we obtained at least 4 million transformants, ensuring at least 40x coverage per variant, estimated from the number of colonies on the plate. Plasmids were isolated the following morning using the Plasmid Plus Midi Kit (QIAGEN).

We transferred the libraries from the mutagenesis plasmid into both the activity plasmid (pCF85) and the DHFR-sandwich abundance PCA plasmid (pCF120). To do this, we digested both the vectors and the mutagenesis-plasmid libraries with the NheI and NotI restriction enzymes (NEB). Digested libraries were purified using a gel extraction kit (QIAGEN). Empty digested vectors were verified by gel electrophoresis and purified using the QIAquick PCR purification kit (QIAGEN). Each linearised vector was assembled with each extracted library using T4 ligase (NEB). The reaction products were dialysed using 0.025 μm mixed cellulose ester (MCE) membranes, concentrated to 5 μl using a SpeedVac machine, and transformed into NEB 10β High-efficiency Electrocompetent *E. coli* cells, followed by 1 h recovery in 2 ml of SOC. Again, 0.1% of the recovery product was plated onto LB agar with ampicillin, and the rest was inoculated into 100 ml of LB liquid with ampicillin. The plasmid was isolated the following morning using the Plasmid Plus Midi Kit (QIAGEN).

To introduce the barcode construct, 2 μg of purified plasmid was digested with the AvrII and BsiWI restriction enzymes (NEB). Digested vectors were verified by gel electrophoresis and purified using the QIAquick PCR purification kit (QIAGEN). The barcode was ordered as a single-stranded oligo from IDT. It was flanked by sequences complementary to those flanking the AvrII and BsiWI sites present in both the activity and abundance plasmids. It was introduced by NEBuilder HiFi DNA Assembly (NEB), following the double-stranded DNA with single-stranded DNA oligo protocol, and transformed into NEB 10β High-efficiency Electrocompetent *E. coli* cells, followed by 1 h recovery in 2 ml of SOC. Again, 0.1% of the recovery product was plated onto LB agar with ampicillin, and the rest was inoculated into 100 ml of LB liquid with ampicillin, then purified using the Plasmid Plus Midi Kit (QIAGEN).

To send the libraries generated for long read sequencing for barcode-variant association, 10 μg of each plasmid was digested with Eco-RV (NEB), The linearized fragments were purified using QIAquick PCR purification kit (QIAGEN). Reads were retained only if both the gene error rate and the barcode error rate (each defined as 1 minus the per-read accuracy) were at or below 1×10⁻⁴ and 1×10⁻³, respectively. Consensus sequences were then called per barcode from the retained reads using alignparse^83^.

### Large-scale transformation and competitions

Each library was transformed in triplicate into *Saccharomyces cerevisiae* BY4741. Three independent precultures were grown in 100 mL YPDA overnight at 30°C. Each replicate was then inoculated into 1 L YPDA at an optical density (OD) of 0.3 and grown for 4 h to an OD of 0.8 to 1.0. Cells were harvested by centrifugation (5 min at 3000 g), washed once in sterile water and once in SORB medium, and resuspended in 43 mL SORB and incubated for 30 min at room temperature (RT) on a rotating wheel. Pre-boiled salmon sperm DNA (875 µL at 10 mg/mL; Agilent Genomics) and 17.5 µg of the corresponding library were added, and they were incubated for a further 10 min. The cells in SORB were combined with 175 mL of plate mixture and shaken for 30 min. DMSO (17.5 mL) was then added, the mixture was split across five 50 mL tubes, and the cells were heat-shocked for 20 min, shaking them every 1.5 min. The tubes were pooled, harvested (5 min at 3000 g), resuspended in 250 mL recovery medium, and allowed to recover for 1 h at 30°C. Cells were harvested again (5 min at 3000 g) and resuspended in 1 L SC-URA. To estimate transformation efficiency, we took 10 µL from the cells in SC-URA and we plated them on selective SC-URA, targeting at least 20-fold library coverage.

For the activity assay, the overnight cell culture was in SC-URA containing 2% raffinose and 0.1% glucose. After 24 h, they were inoculated into 1 L of the same medium at an OD of 0.3. The following day, cells were inoculated into 1 L SC-URA containing 2% galactose and 0.1% glucose at an OD of 0.05 to induce kinase overexpression. The remaining cells were harvested (10 min at 3000 g) and frozen for DNA extraction (input). Cells grown in galactose were harvested the next day at an OD of 1.6 (10 min at 3000 g) and frozen for DNA extraction (output).

For the stability assay, the overnight SC-URA culture was inoculated the following day into SC-URA-ADE at an OD of 0.3. The next day, cells were inoculated into 1 L SC-URA-ADE+methotrexate (200 µg/mL) to select for cells expressing stable kinase variants. The remaining cells were harvested (10 min at 3000 g) and frozen for DNA extraction (input).Cells grown in SC-URA-ADE+methotrexate were harvested at an OD of 1.6 (10 min at 3000 g) the next day and frozen for DNA extraction.

### DNA extraction, plasmid quantification and library preparation

DNA was extracted from a total number of harvested cells equivalent to 500 mL of culture at an OD of 1.6, following the protocol described previously in Faure et al.^25^ and Weng et al.^26^ Plasmid concentration in the extracted samples was quantified in triplicate by quantitative PCR (qPCR) against a standard curve of known concentrations, using oGJJ152 and oGJJ153 as primers, which amplify the origin of replication common to both the toxicity and sPCA assay plasmids.

To prepare the barcode libraries for Illumina sequencing, we performed two subsequent rounds of PCR amplification. In the first PCR (PCR1), we amplified the inserted barcode from each sample while adding the Illumina adapters and increasing nucleotide complexity by introducing frameshift primers. PCR1 was performed independently for each sample with a common set of frameshift primers, as the regions flanking the barcode are identical across libraries and plasmids. A minimum of 100 million copies of the extracted plasmid was used as starting template for each sample. 10 PCR1 cycles were performed using Q5 Hot Start High-Fidelity DNA Polymerase (NEB). The reactions were cleaned using 2 µL of ExoSAP per 50 µL reaction and then column-purified with a MinElute PCR Purification Kit (QIAGEN). The purified product was used as starting template for the second PCR (PCR2), amplified with the standard i5 and i7 Illumina primers (dual indexing) for 10 cycles. PCR2 products were run on a 2% agarose gel for quantification and pooled at equimolar ratios. The final pooled library was run again on a 2% agarose gel and purified by gel extraction using the QIAEX II Gel Extraction Kit (QIAGEN). The final product was subjected to Illumina paired-end 2×50 bp sequencing on a NovaSeq 6000 instrument.

### Sequencing data processing and thermodynamic modeling

FastQ files from every sPCA and toxicity experiment were converted to variant fitness estimates with DiMSum^82^, run under default parameters apart from two adjustments. First, the “barcodeIdentityPath” argument was given the barcode-variant table from the PacBio run, so that counting was confined to the genotypes present in our SUNi-derived library and reads arising from sequencing error were discarded. Second, a per-variant threshold on input reads for 1-nt changes was applied via “fitnessMinInputCountAny”, to suppress the contribution of sequencing error to lower-order mutants.

Biophysical parameters were inferred with MoCHI^43^, taking the raw single- and double-mutant fitness values as input. For each kinase we fitted a separate phenomenological enzyme folding and activation model, which assumes the protein exists in three states: unfolded and inactive; folded and inactive; and folded and active, as described in Beltran et al. (ref). The model estimates the single-mutation Gibbs free energy effects on folding (ΔGf) and activity (ΔGa), under the assumption that the free energy changes of variants (ΔΔGf and ΔΔGa) are additive. The network was configured with two additive-trait layers, encoding folding and activation energies, together with one linear layer for each assay (activity and sPCA). We used the non-linear transformations TwoStateFractionFolded and ThreeStateFractionBound for the abundance (sPCA) and activity readouts, respectively, derived from the Boltzmann distribution, which relates the proportion of folded molecules, and of molecules in the active versus inactive state, to their free energies. Cross-validation accuracy (Pearson r between measured and predicted double-mutant fitness) was corrected for attenuation as r_corrected = r_observed / √(R_xx · R_yy), where R_xx is the reliability of the measurements, taken for each kinase and assay as the median pairwise correlation of the three fitness replicates (predictions were treated as error-free), and R_yy is the reliability of the predictions, which we set to 1.

### Protein structures, structure metrics, and contacts

The five kinase domain sequences were aligned according to the KinCore multiple sequence alignment of the kinase domain^84^, which defines the shared aligned positions (Pos_al) used for all cross-kinase comparisons, region assignments and functional annotations (Table S3)

In the same way, conserved elements of the kinase domain were annotated from the region definitions of the KinCore alignment^84^: every position was labelled with its structural element, the β-strands (β1-β5), the α-helices (αC to αI), the catalytic loop, activation loop, and the F-loop. Each residue was assigned to the N-lobe, hinge or C-lobe by its position in the alignment. All annotations, including the C- and R-spines, the glycine-rich loop, the gatekeeper, the DFG/Mg-positioning motif, the HRD catalytic residues and the substrate-positioning loop, were annotated per aligned position.

Structural models of the five kinase domains were generated with the AlphaFold3 web server^85^, providing the kinase-domain sequence segment covered by the mutagenesis library and including ATP as a bound ligand in each run. All structural metrics were derived from these ATP-bound models.

For every residue we computed the relative solvent-accessible surface area (rSASA) with the freeSASA package (v2.2.1)^86^. Residues with rSASA < 0.25 were assigned to the core and the rest to the surface.

Pairwise residue distances within each kinase were defined as the shortest distance between any two side-chain heavy atoms of the residue pair. Residue-to-ATP distances were obtained in the same way, as the minimum heavy-atom separation between each residue and the ATP ligand. We additionally computed the active-site distance, defined as the minimum heavy-atom distance from each residue to either the ATP ligand or the catalytic aspartate of the HRD motif (alignment position 161, an aspartate in all five kinases).

Residue contacts were calculated on each ATP-bound model with getContacts (https://getcontacts.github.io/). Residues making one or more contacts with ATP were flagged as ATP-contacting. The active site was then defined as the union of these ATP-contacting residues and the residues annotated as substrate-positioning or Mg-positioning.

### Identification of allosteric activating, inactivating, stabilizing and destabilizing mutations, and regional enrichments

To compare mutational effects across the five kinases, the inferred ΔΔGf and ΔΔGa values and their errors were first normalised to a common scale using reference mutations expected to be strongly affected in every kinase (αF-helix proline substitutions for folding and Mg-positioning active-site substitutions for activity).

The normalised mean weights and standard errors from the MoCHI fits were then used to identify mutations with significant effects on folding or activity by z-test, with P values derived from a normal distribution and the false discovery rate controlled across the pooled variants of the five kinases by the Benjamini-Hochberg procedure (FDR < 0.1). Using an effect-size threshold of 0.2 kcal/mol, variants were classified as stabilizing or destabilizing and as activating or inactivating, and activating or inactivating mutations lying outside the active site were considered allosteric. Enrichment of a given class within secondary structure elements, functional regions or individual sites was assessed by Fisher’s exact test, comparing against the rest of the kinase domain as background. To define ‘major allosteric sites’, each position outside the active site was tested separately for an excess of activating and of inactivating mutations relative to the rest of the same kinase, using a one-sided Fisher’s exact test (Benjamini-Hochberg FDR < 0.1); positions enriched for activating mutations were taken as activating allosteric sites, those enriched for inactivating mutations as inhibitory allosteric sites.

### Quantification of the distance dependence of mutational effects

To quantify how activity effects depend on distance from the active site, we defined for each residue its distance to the active site as the minimum heavy-atom distance to either the bound ATP or the catalytic aspartate of the HRD motif, taking the smaller of the two. For each kinase we fitted an exponential curve |ΔΔGa| = a·e^k·d^, where *d* is the distance to the active site, *a* is the effect size at the active site (|ΔΔGa|₀) and *k* is the decay rate, using nls() in R with starting values from optim(). The half-distance d½ = ln(2)/|k| summarised the range over which effects decay. Fits were carried out on all mutations, on the per-residue median |ΔΔGa|, and separately on activating (ΔΔGa < 0) and inactivating (ΔΔGa > 0) mutations. We also fit the curve across secondary structure types, with residues grouped as α-helix, β-strand or loop from their assigned region.

To quantify the dependence of activity effects on spatial direction, we measured the decay separately along six directions defined in the reference frame of the 2SRC ^56^ crystal structure of inactive Src. Each kinase structure was superposed onto 2SRC so that all Cα coordinates shared the 2SRC axes, and the active site was taken as origin using the heavy atoms of the bound nucleotide and of the catalytic HRD aspartate. For each residue we computed its signed displacement along the x, y and z axes, and for each of the six half-axes (x±, y±, z±) we retained residues lying in that direction and within 10 Å of the origin along the two orthogonal axes. The same exponential decay |ΔΔGa| = a·e^k·d^ was fitted along each direction, with *d* the displacement along the chosen axis, and the half-distances d½ = ln(2)/|k| were compared across directions and kinases.

### Analysis of surface pockets

Surface pockets were mapped for each kinase with FTMap^68^. Following the approach of the Kinase Atlas^69^, each AlphaFold3 structure was split at aligned position 113 into its N- and C-terminal lobes, which were mapped separately so as to reduce the dominance of the ATP-binding pocket increase the relative weight of surface pockets, and the bound ATP was removed. Each resulting cross-cluster of docked probe molecules, was defined as a consensus site, and was taken as one candidate pocket. The druggability score for each pocket was defined as the number of probe clusters in the site, and the residues forming the pocket as the protein residues contacting the docked probes.

To obtain a set of pockets comparable across the five kinases, all candidate pockets were pooled and defined in the rest of kinases by their residues in aligned-positions. For each pair of pockets we computed the Szymkiewicz-Simpson overlap coefficient which is the size of the intersection divided by the size of the smaller residue set, converted it to a distance (*1-coefficient*), and clustered the pockets by hierarchical clustering, cutting the tree at a height of 0.6 to define global pockets. Global pockets were numbered in order of their most N-terminal position, so that the pocket containing the most N-terminal residue is pocket 1.

For each global pocket in each kinase we summarised the activity effects of its residues (median per-residue ΔΔGa) and its druggability, and tested whether allosteric activating or inactivating mutations were over-represented among its residues. Enrichment was assessed for each pocket against the whole kinase as background with a hypergeometric test (Benjamini-Hochberg FDR < 0.1) and reported as an odds ratio. For testing specific-kinase pockets, for each kinase we subtract its whole-domain (non-active-site) median activity ΔΔGa to remove the kinase-level baseline offset, then within each pocket we compare every kinase pair on these baseline-corrected values with a two-sided Wilcoxon rank-sum test, reporting Cliff’s delta as the effect size and BH-adjusting the p-values (FDR < 0.1).

## Supplementary Figure Legends

**Figure S1. Experimental design, reproducibility, and per-position fitness.** (A) Single-substitution fitness heatmaps for the N-terminal region of the kinase domain measured in the pilot activity-dependent toxicity assay, for the four kinases tested (FGR, ZAK, JNK2, TSSK2); rows, mutant amino acid; columns, position. (B) Correlation between the three replicates (R1 to R3) of the activity and abundance assays for FGR, ZAK, JNK2 and TSSK2 (Pearson’s R). (C) Fitness distributions of single and double mutants for the abundance and activity assays of the same four kinases; dashed lines mark the mean of stop codons (red) and synonymous variants (green). (D) Normalised fitness of single substitutions at ATP-contact versus non-ATP-contact positions in the pilot toxicity assay, per kinase. (E) For each single substitution, the number of distinct double-mutant backgrounds in which it was measured (1 to 8), per kinase and assay (S, abundance; T, activity). (F) Median abundance fitness per aligned position for the five kinases. (G) As in (F) for median activity fitness. (H) Schematic of the competition selection: a pooled library of yeast expressing variants (input) is selected and sequenced before and after selection (output). (I) Domain architecture of the five full-length kinases.

**Figure S2. MoCHI evaluation.** (A) Observed against MoCHI-predicted fitness in the abundance assay, per kinase (Pearson’s r). (B) As in (A) for the activity assay. (C) Observed abundance fitness as a function of ΔΔGf, per kinase; red line, model fit.

**Figure S3. Energetic maps of folding and activity.** (A) ΔΔGf of every single amino acid substitution in each kinase. Rows are mutant amino acid and columns are aligned positions, with the wild-type sequence shown below each map. Unmeasured substitutions are left blank. (B) As in (A) for ΔΔGa.

**Figure S4. Distributions, structural context, and cross-kinase comparison of folding energetics.** (A) Distribution of ΔΔGf per kinase. ΔΔGf = 0 as dashed line. (B) Median ΔΔGf per aligned position for the five kinases. Letters are the wild-type residue. The panel below marks secondary structure and functional regions. (C) ΔΔGf distributions of core versus surface positions per kinase. The OR is the enrichment of destabilising mutations in the core relative to the surface (Fisher’s exact test); ****, p < 0.001. (D) Pairwise Pearson correlation of per-position median ΔΔGf between kinases, computed separately within each secondary structure region. (E) Pairwise scatter of per-position median ΔΔGf between kinases (Pearson’s R correlation). (F) Percentage of destabilising mutations in the N-lobe (dark pink) and C-lobe (light pink) of each kinase. Odds ratio for the enrichment of destabilising mutations in that lobe versus the rest of the domain is written above each bar. Per-kinase Fisher’s Exact Test (FET) comparing the two lobes is shown (*** FDR < 0.001, ns not significant). (G) Enrichment of destabilising and stabilising mutations within secondary structure elements (columns) for each kinase (rows) (FET). Red asterisks are enrichment for destabilising mutations (FDR < 0.1); white asterisks are enrichment for stabilising mutations (FDR < 0.1). AL, activation loop; CL, catalytic loop; FL, F loop. (H) Close-up of the αF helix in each kinase, coloured by the per-position median deviation of folding free energy from the cross-kinase median (ΔΔΔGf = kinase ΔΔGf - cross-kinase median ΔΔGf for the same substitution). (I) As in (E), restricted to αF positions (215 to 237); diagonal line, y = x. (J) Median deviation of ΔΔGf from the cross-kinase consensus (median ΔΔΔGf) at αF positions. Letters are the wild-type residue. Asterisks mark when ΔΔΔGf deviates from zero at FDR < 0.1 (one-sample Wilcoxon signed-rank test).

**Figure S5. Activity effects at the active site**. (A) Active-site positions shown as spheres on each kinase structure, coloured by median ΔΔGa. ATP shown as yellow sticks. (B) Enrichment of inactivating and activating mutations in the active site relative to the rest of the domain, per kinase and pooled across the five kinases (Fisher’s exact test; ***, FDR < 0.0001; **, FDR < 0.001; *, FDR < 0.1).

**Figure S6. Distance decay of allosteric effects and enrichment of allosteric mutations.** (A) Top row, per-residue median |ΔΔGa| versus distance to the active site, per kinase. Points are coloured by ATP direct contact (yellow), or other (grey), with error bars showing the standard error of the median and a black exponential decay fit. Bottom row, the same for all single mutations. (B) Per-residue median |ΔΔGa| versus distance to the active site fitted separately within each secondary structure type, per kinase. (C) Enrichment of activating mutations in the N-lobe versus C-lobe per kinase. Each bar is the log2 odds ratio of the enriched lobe against the other lobe (FET,* p < 0.05, ** p < 0.01, *** p < 0.001). (D) Enrichment of inhibitory allosteric mutations within secondary structure regions (rows) per kinase (columns) (FET, asterisks FDR < 0.1). (E) Pooled log2 odds ratio enrichment of each amino acid among allosteric activating (top) and allosteric inactivating (bottom) mutations outside the active site, by wild-type amino acid (left) and mutant amino acid (right). Bars are coloured by amino-acid type (FET, asterisks FDR<0.1). (F) As in (D) for activating allosteric mutations. (G) Number of distinct activating allosteric substitutions that activate 1 to 5 kinases. (H) As in (E), per kinase, for activating (top, purple) and inhibitory (bottom, red) allosteric mutations; grey, fewer than three mutations (Fisher’s exact test; asterisks, enrichment at FDR < 0.1).

**Figure S7. Conservation of major allosteric sites.** (A) Median ΔΔGa per aligned position for the five kinases. Letters are wild-type residues. Red outlines mark positions enriched for inhibitory allosteric mutations and blue outlines positions enriched for activating allosteric mutations. Grey shading marks active-site residues. The number above each column is the number of kinases in which the position is a major allosteric site. The track below marks secondary structure and functional regions. (B) Surface enrichment of hotspots as a function of the number of kinases sharing the hotspot (FET, asterisks p < 0.5). (C) Per-residue cross-kinase Spearman correlation (ρ) of ΔΔGa for core versus surface positions.

**Figure S8. Structural, sequence, and functional conservation and divergence of pockets.** (A) FTMap probe clusters (blue) on the SRC, FGR, JNK2 and TSSK2 structures. ATP is shown as yellow sticks. (B) Per-pocket functional conservation, defined as the mean cross-kinase correlation of the ΔΔGa effects of the pocket’s residues (“effect r”), versus structural divergence (mean pairwise Cα RMSD; top) and versus sequence conservation (mean residue identity; bottom). Each point is a pocket, labelled by pocket number and coloured by its median ΔΔGa. Dashed lines mark the means. (C) Per-position activity effects and directional allosteric enrichment across the 37 structural pockets. Each panel is one global pocket. Within a panel, rows are the five kinases and columns are the pocket’s aligned residue positions. Tile colour is the per-position median ΔΔGa, white tiles mark positions not present in that kinase. The two rightmost columns give the enrichment of activating (A) and inhibitory (I) allosteric mutations in the pocket in each kinase; asterisks, enrichment at FDR < 0.1 (one-sided hypergeometric test). (D) Pairwise comparison of pocket activity effects between kinases. Cell colour is Cliff’s delta for the row kinase relative to the column kinase, comparing ΔΔGa of substitutions at the pocket’s non-active-site residues (blue, row shifted toward activation; red, toward inactivation). An asterisk marks pairs differing at FDR < 0.1 (two-sided Wilcoxon rank-sum test).

**Figure S9. Pocket array.** ChimeraX plots of the 37 pockets mapped onto the aligned structures of the five kinases, showing the location of each pocket in each kinase.

## Supplementary Tables

**Table S1. Variant fitness.** Fitness scores for all variants assayed across the five kinases in the activity (toxicity) and abundance (sPCA) assays. Columns are the kinase, assay, amino acid sequence, number of amino acid changes, wild-type and stop-codon flags, the fitness estimate and its standard error, and the per-replicate values (replicates 1 to 3).

**Table S2. All ΔΔG.** Normalised ΔΔG (kcal/mol) with standard error for every single substitution in the activity and folding models across the five kinases. Columns are the kinase, assay, domain and aligned position, wild-type and mutant residue, and structural annotations (relative solvent accessibility, core/surface, secondary structure, region, lobe, ATP contact, active site, functional feature).

**Table S3. Kinase domain alignment.** KinCore-based alignment of the five kinase domains. Each row is an aligned position giving the structural region and the residue of each kinase. A dash represents an alignment gap.

**Table S4. Pocket properties.** The 37 pockets, one row per pocket and kinase. Each row gives the pocket number and classic name, predicted inhibitor type, the kinase’s member residues and their count, the pocket size, the median activity effect (kcal/mol), the druggability score, and the directional allosteric enrichment of activating and inactivating mutations (odds ratio and FDR).

## Supplementary Movies

AlphaFold model of each kinase domain shown with their surface map, coloured by the per-position median ΔΔGa. The bound ATP is shown in yellow. Each movie rotates the structure 360° about the vertical axis. Movie 1, SRC; Movie 2, FGR; Movie 3, ZAK; Movie 4, JNK2; Movie 5, TSSK2.

Supplementary Video 1 - SRC median ΔΔGa surface map

Supplementary Video 2 - FGR median ΔΔGa surface map

Supplementary Video 3 - ZAK median ΔΔGa surface map

Supplementary Video 4 - JNK2 median ΔΔGa surface map

Supplementary Video 5 - TSSK2 median ΔΔGa surface map

## Acknowledgements

This work was supported by Wellcome (Grant reference: 220540/Z/20/A,‘Wellcome Sanger Institute Quinquennial Review 2021-2026’), a European Research Council (ERC) Advanced (883742) grant, the Spanish Ministry of Science and Innovation (LCF/PR/HR21/52410004, EMBL Partnership, Severo Ochoa Center of Excellence), Agència de Gestió d’Ajuts Universitaris i de Recerca (AGAUR, 2021 SGR01226), and the CERCA Program/Generalitat de Catalunya. A.B. was supported by a La Caixa Junior Leader Fellowship (LCF/BQ/PR25/12110006).

## Author contributions

C.F. performed all experiments and analyses. C.F., A.B. and B.L. conceived the project, designed analyses, and wrote the manuscript.

## Data availability

All sequencing data have been deposited in the European Nucleotide Archive (ENA) at EMBL-EBI under accession number PRJEB122867.

## Code availability

All scripts used in this study are available at: https://github.com/lehner-lab/KINASE-MAPS

