## Supplementary figures S1 to S8 for "Functional pocketomes and complete allosteric maps of five protein kinases"

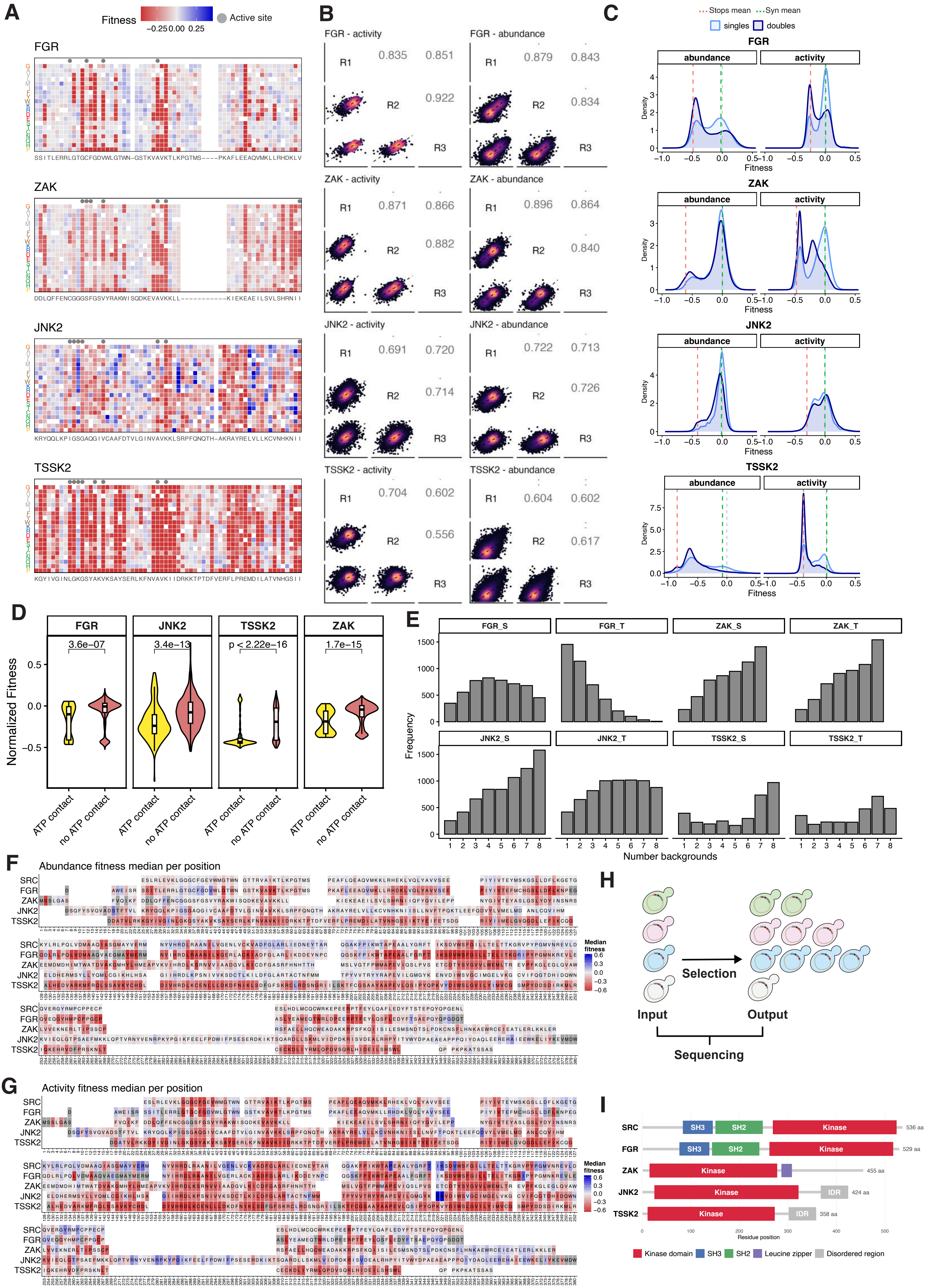


**Supplementary Figure 1**

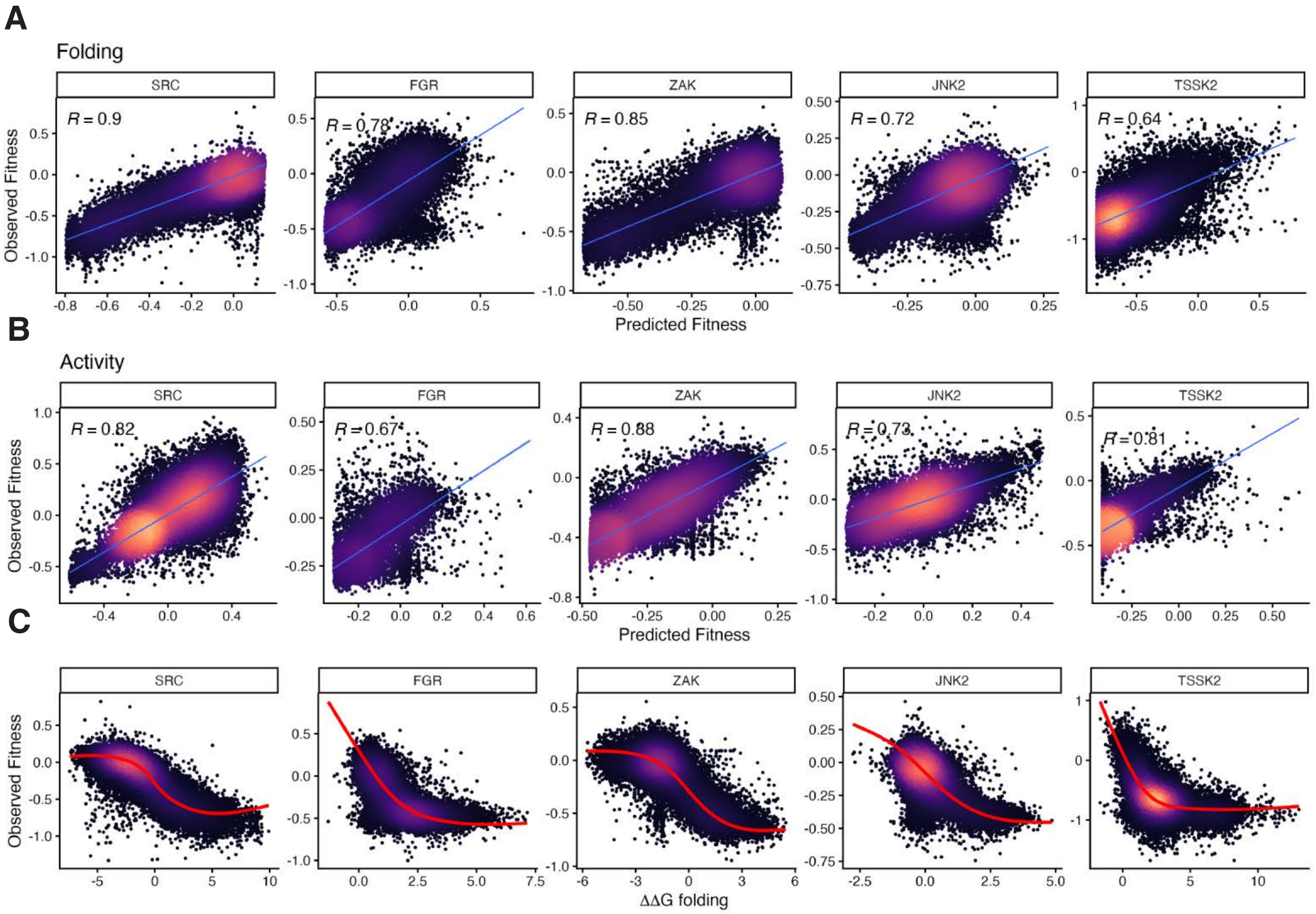


**Supplementary Figure 2**

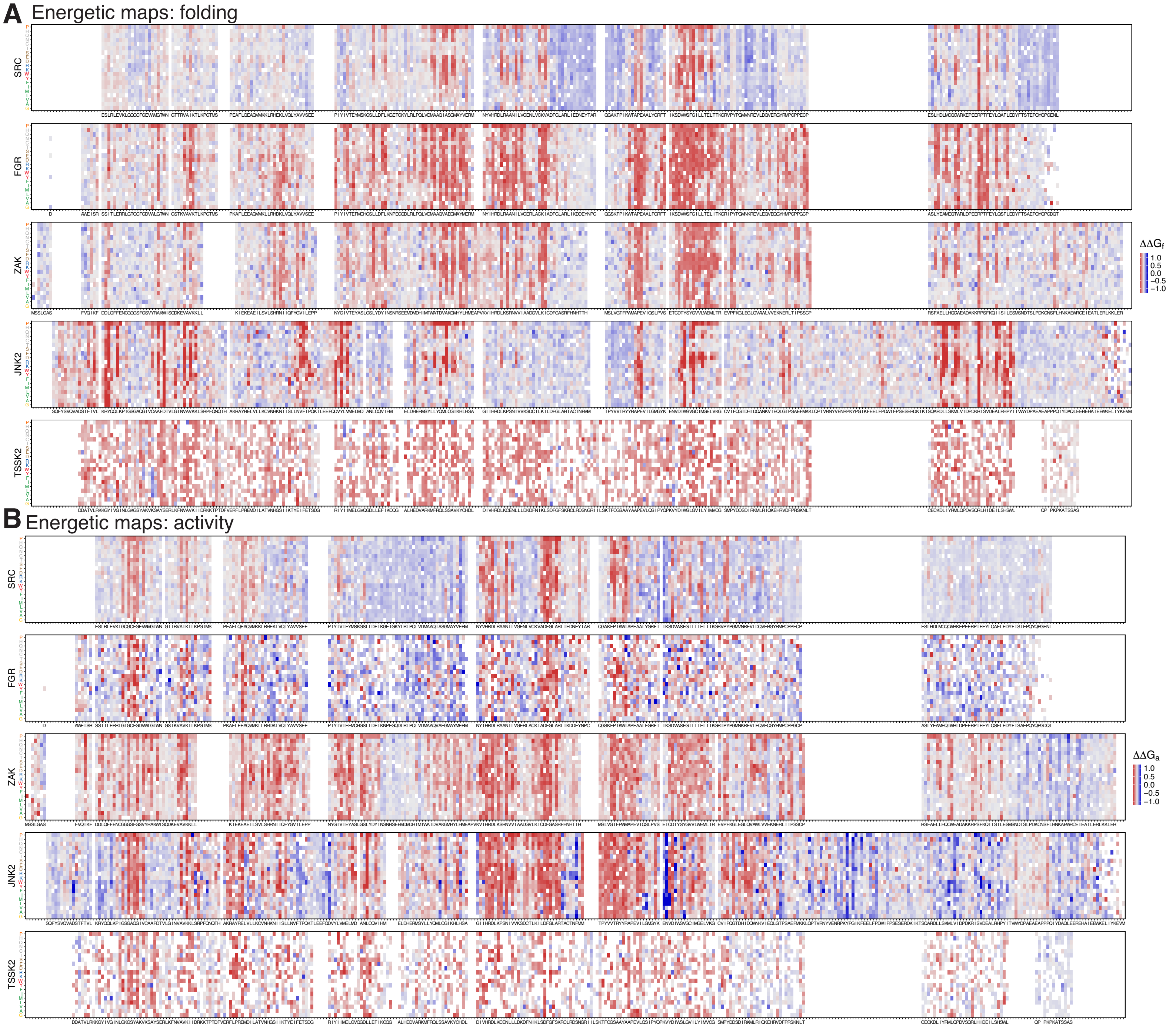


**Supplementary Figure 3**

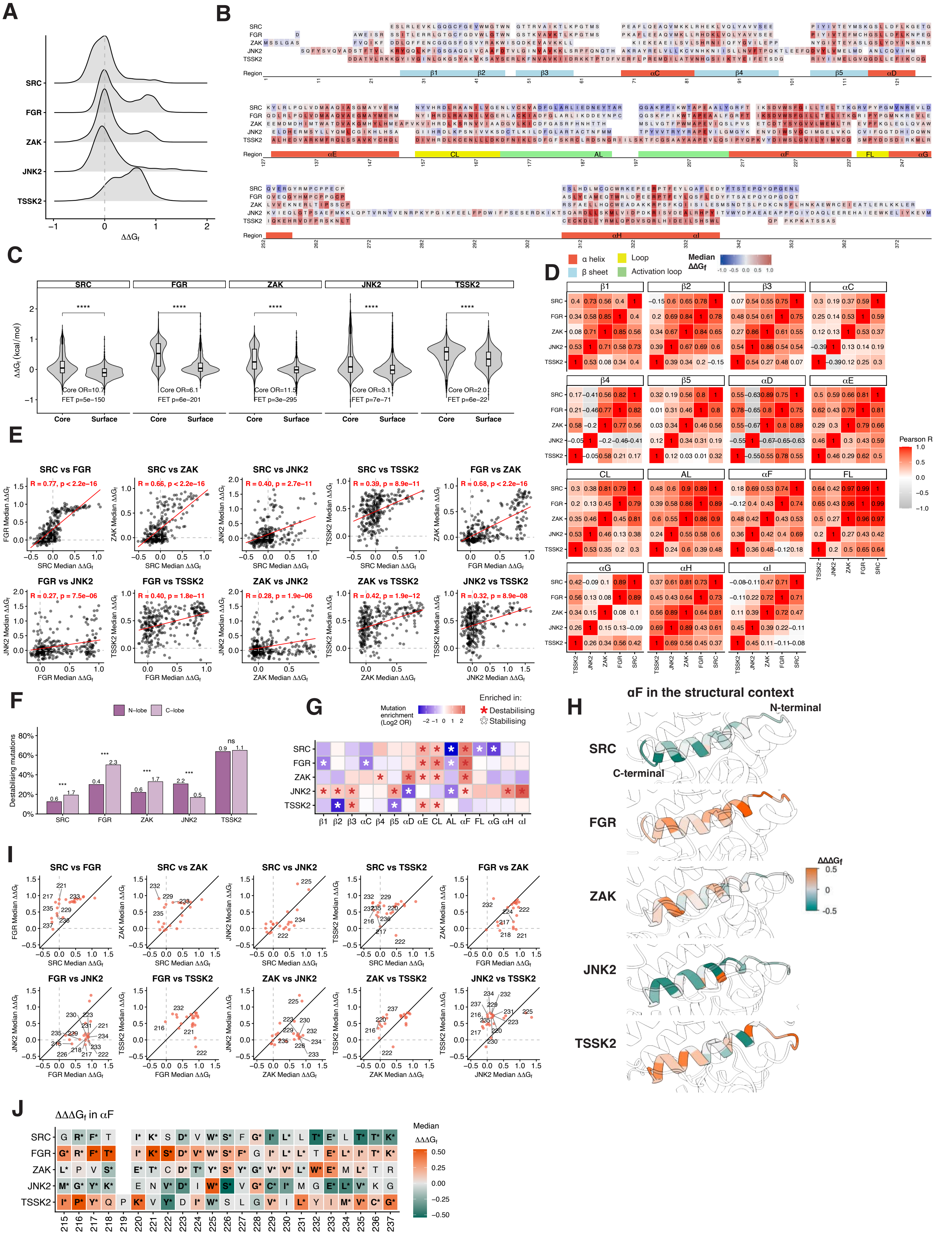


Supplementary Figure 4

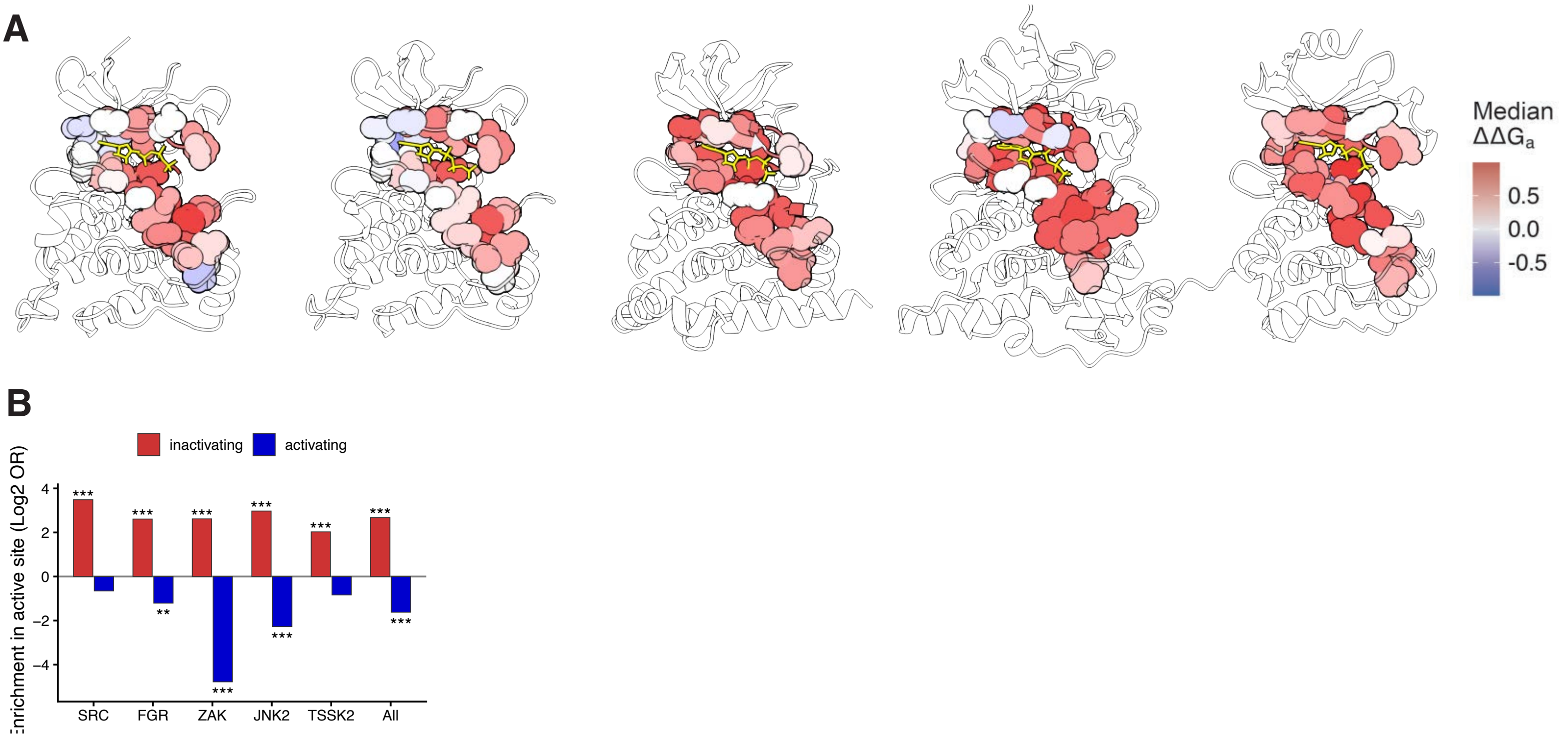


**Supplementary Figure 5**

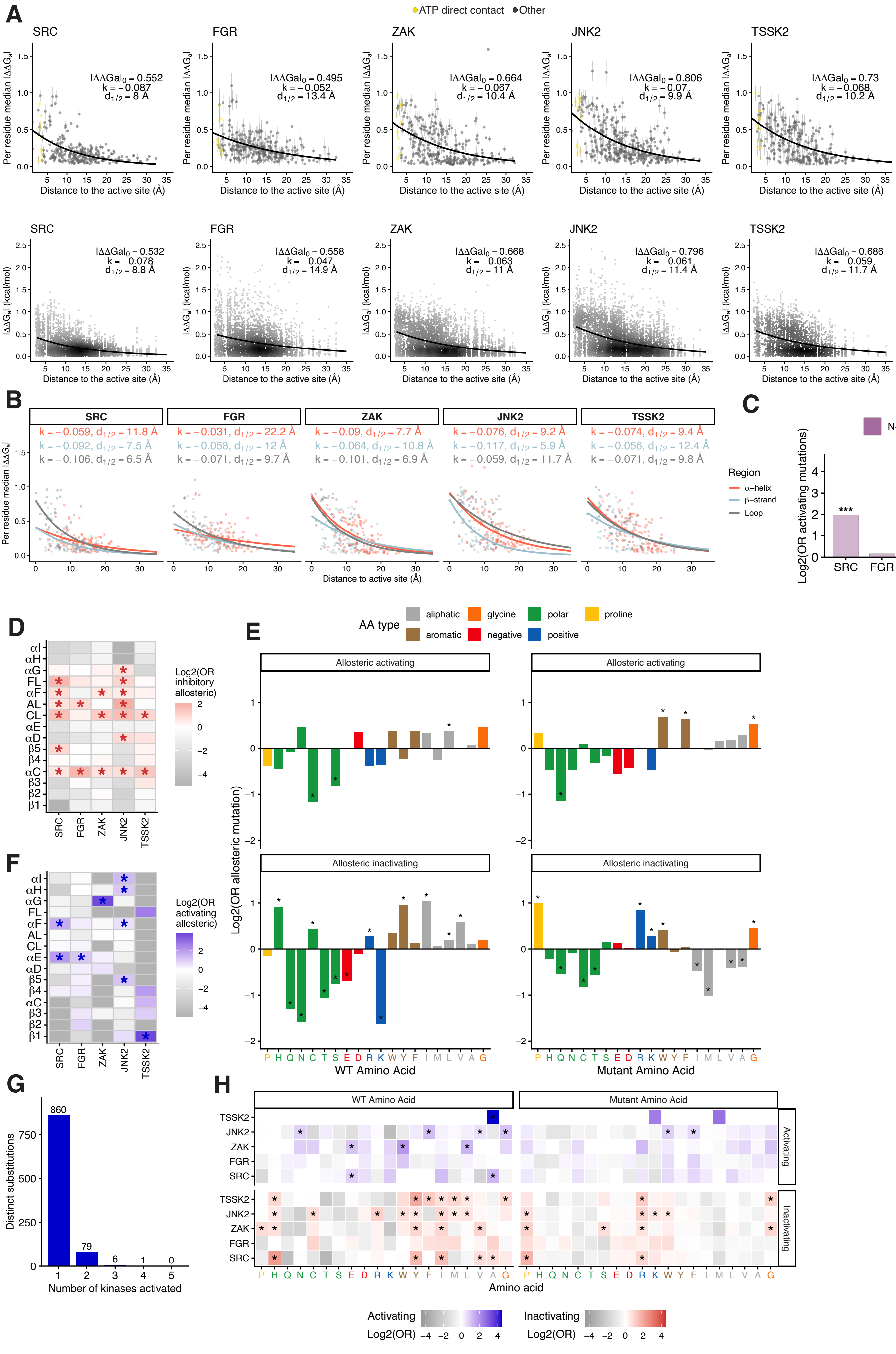


**Supplementary Figure 6**

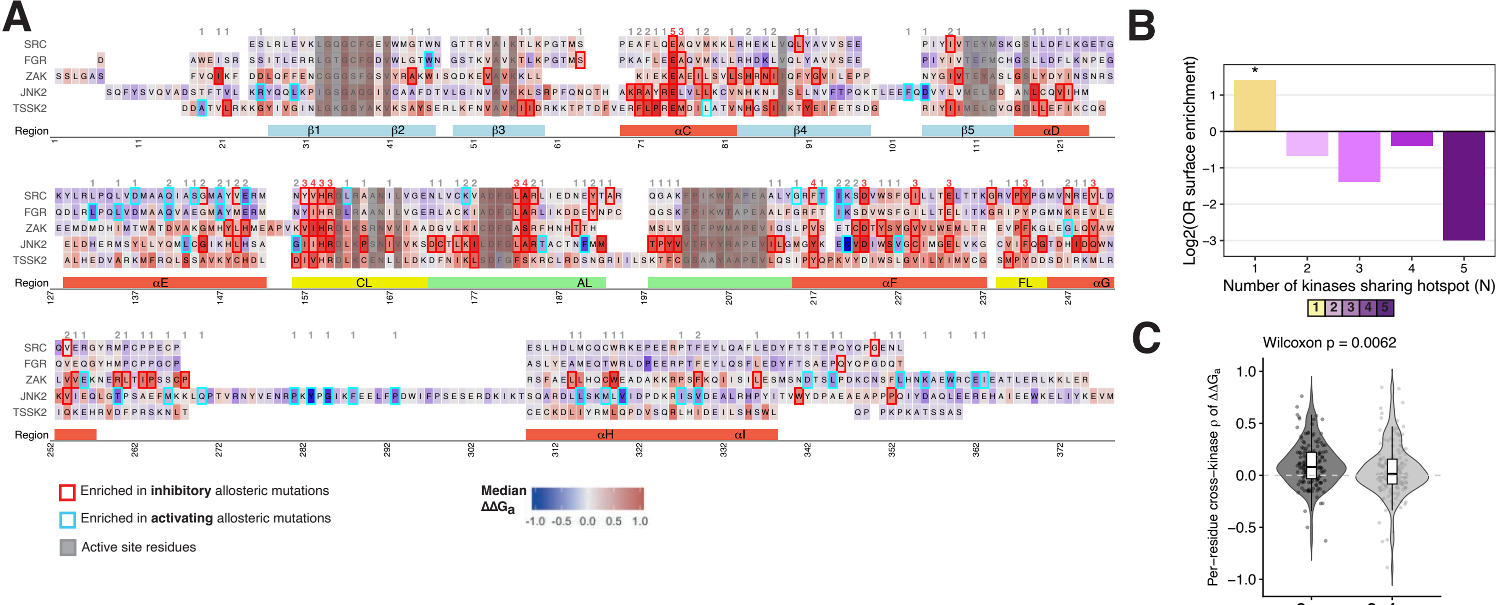


### Supplementary Figure 7

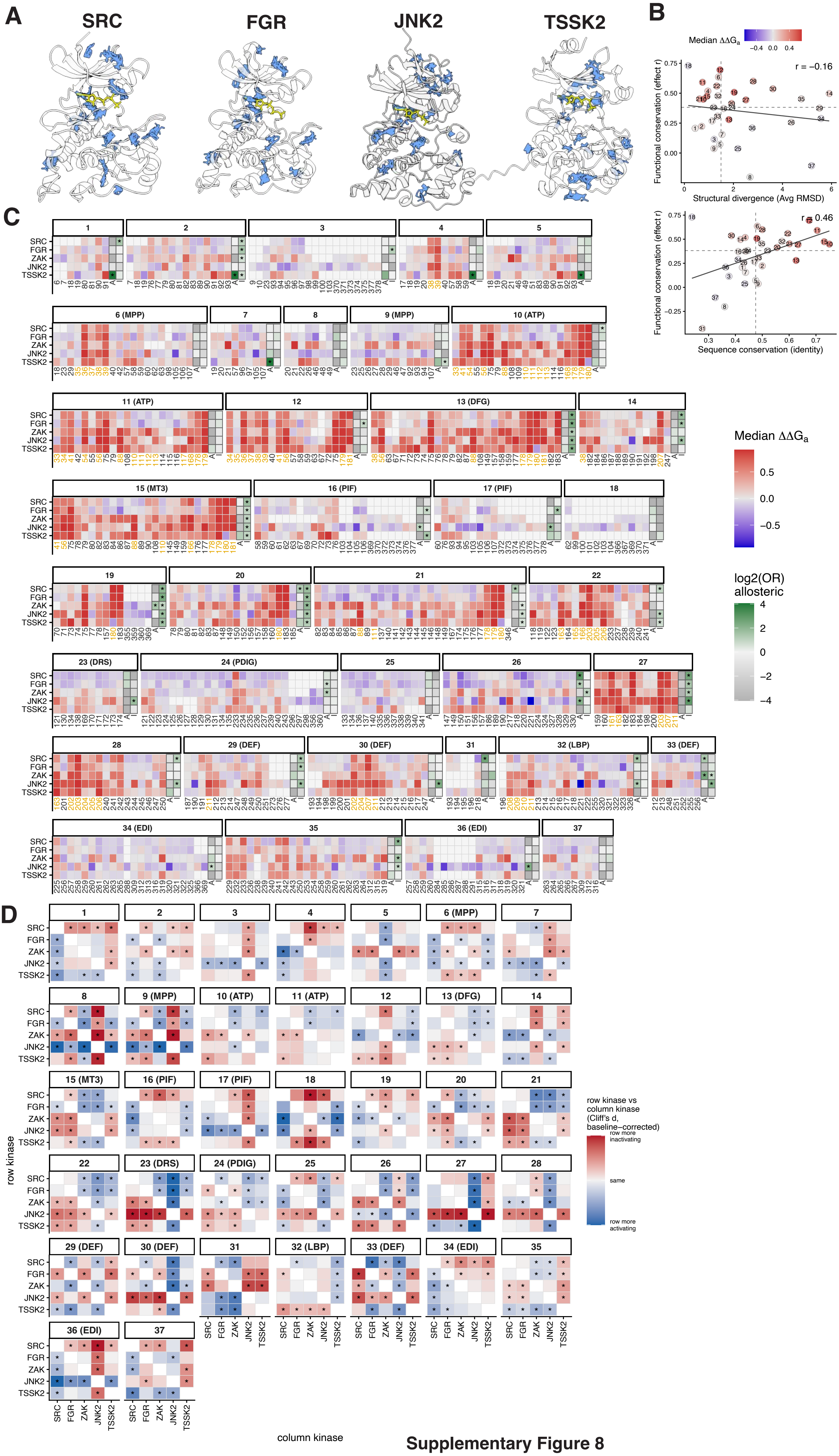


Supplementary Figure 8
