## Supplementary figure S9 for "Functional pocketomes and complete allosteric maps of five protein kinases"

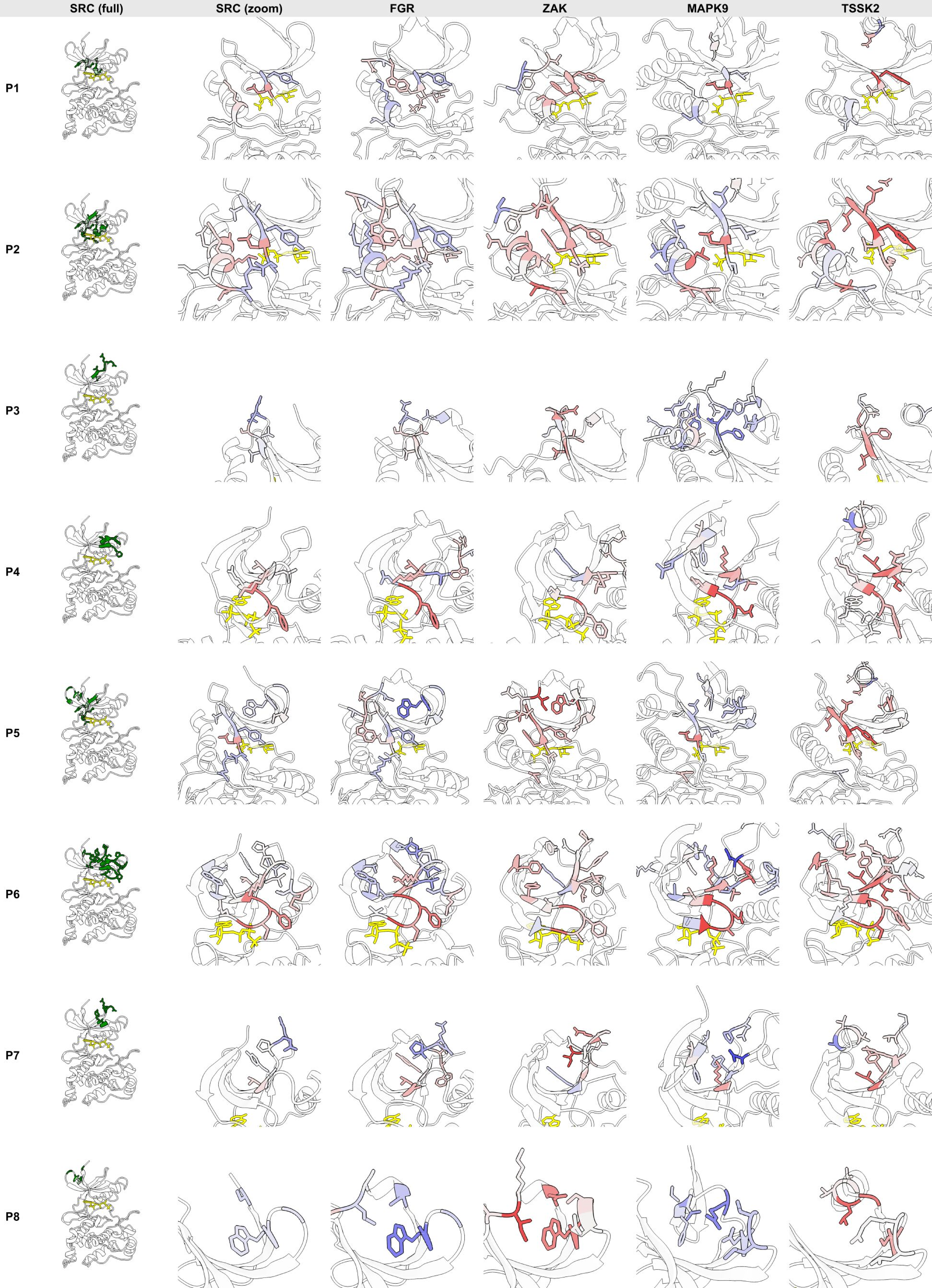
SRC (full)
SRC (zoom)
FGR
ZAK
MAPK9
TSSK2
P1
P2
P3
P4
P5
P6
P7
P8

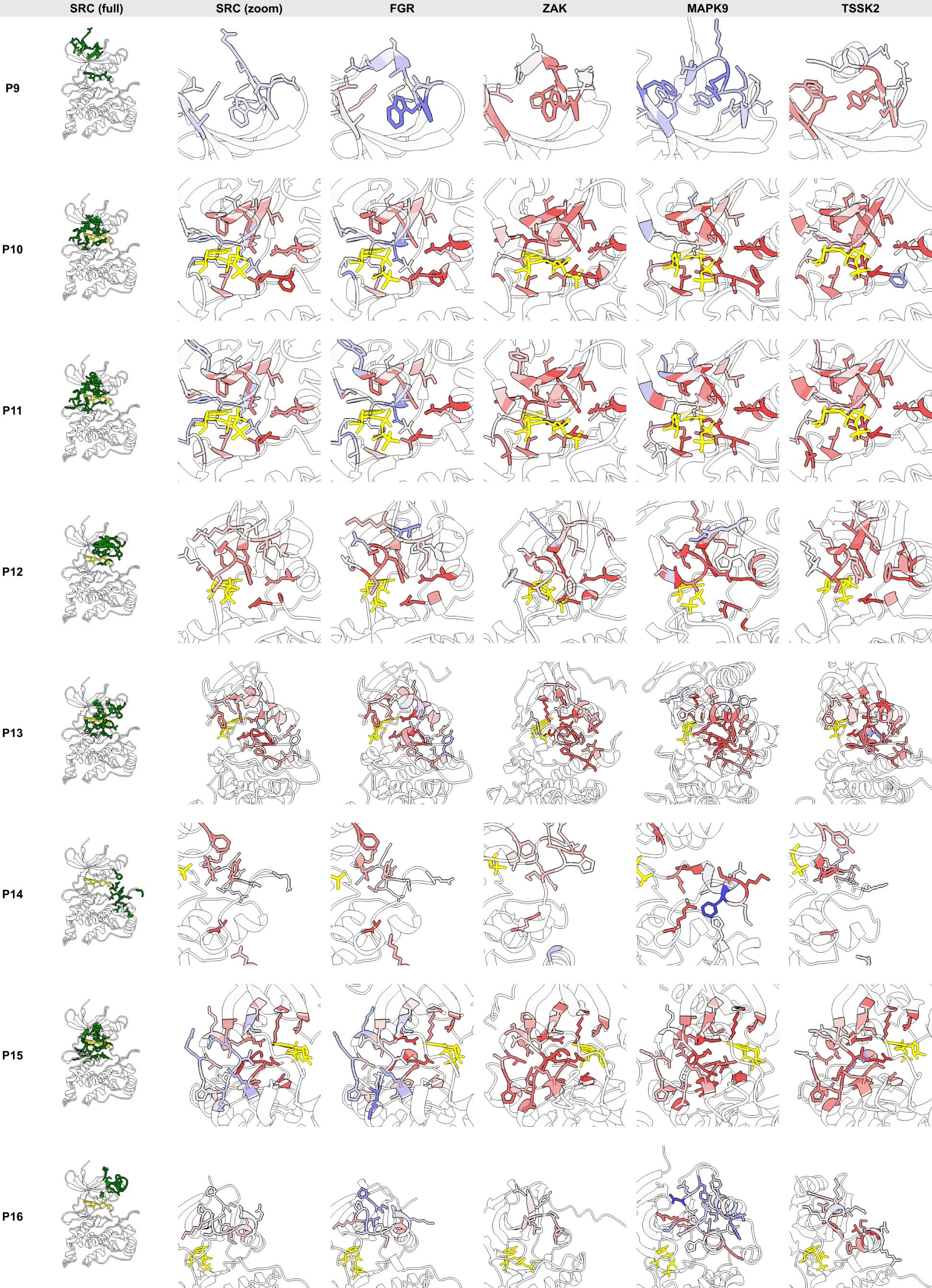

SRC (full)
SRC (zoom)
FGR
ZAK
MAPK9
TSSK2
P9
P10
P11
P12
P13
P14
P15
P16

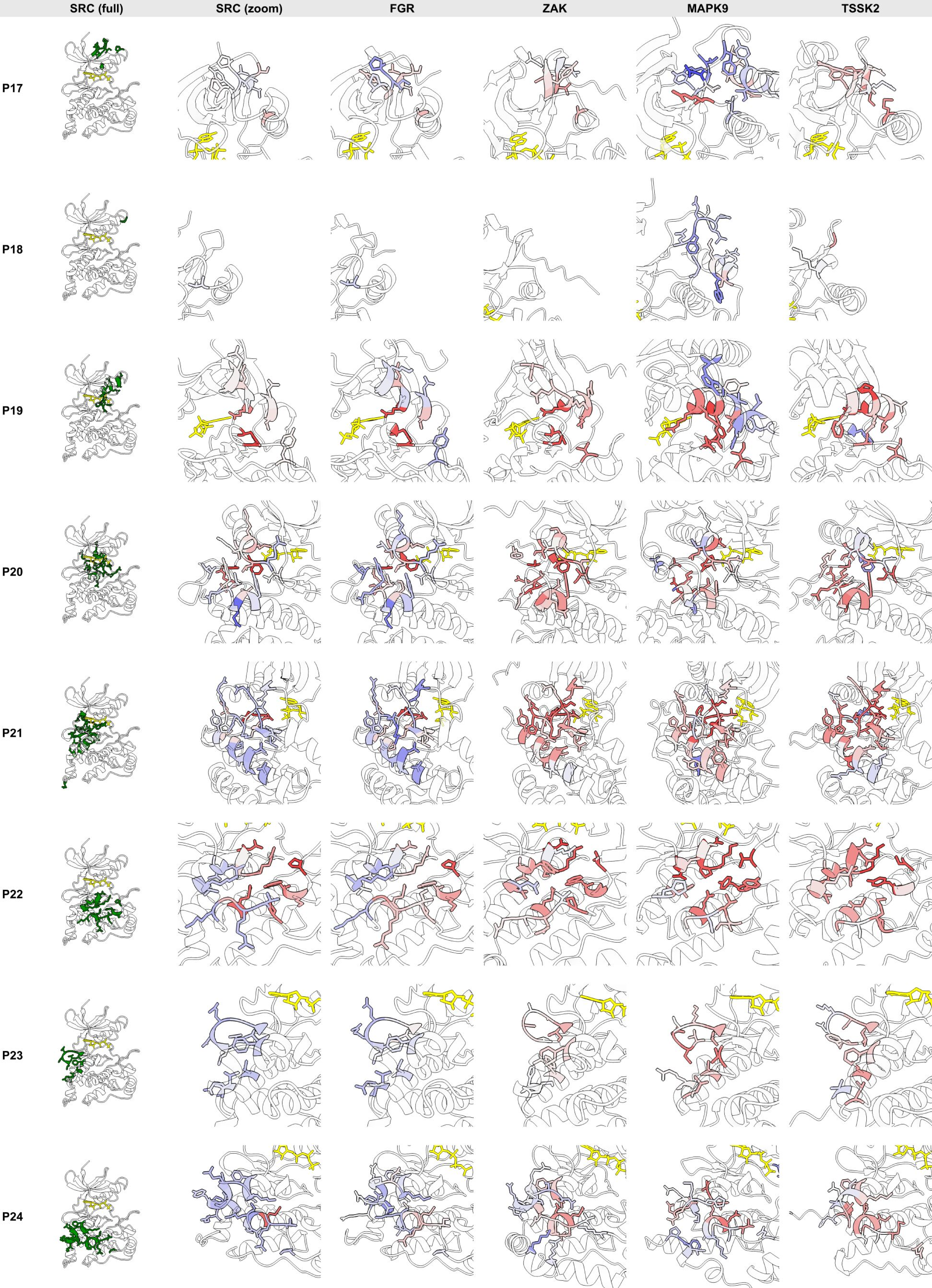

SRC (full)
SRC (zoom)
FGR
ZAK
MAPK9
TSSK2
P17
P18
P19
P20
P21
P22
P23
P24

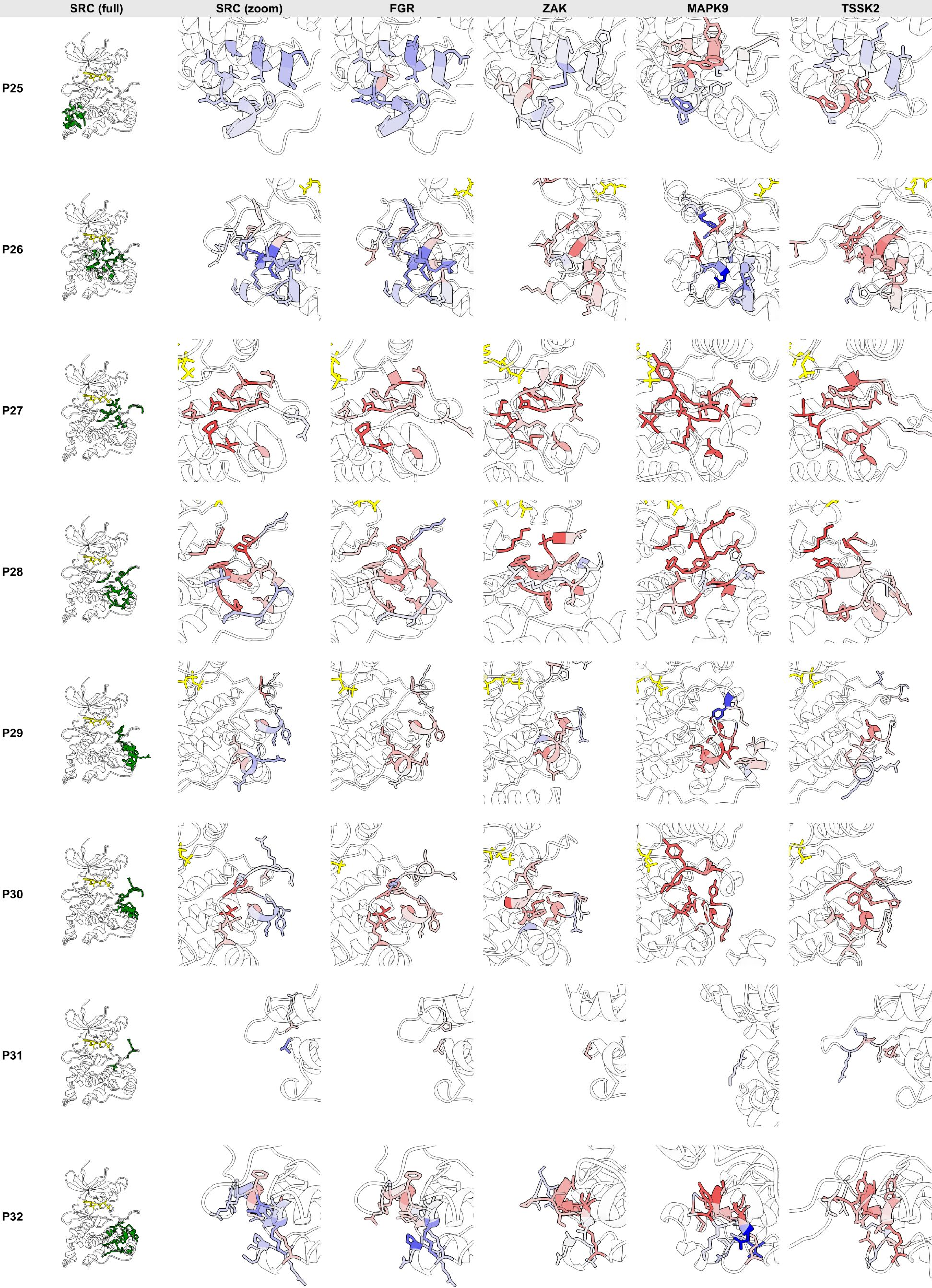
SRC (full)
SRC (zoom)
FGR
ZAK
MAPK9
TSSK2
P25
P26
P27
P28
P29
P30
P31
P32

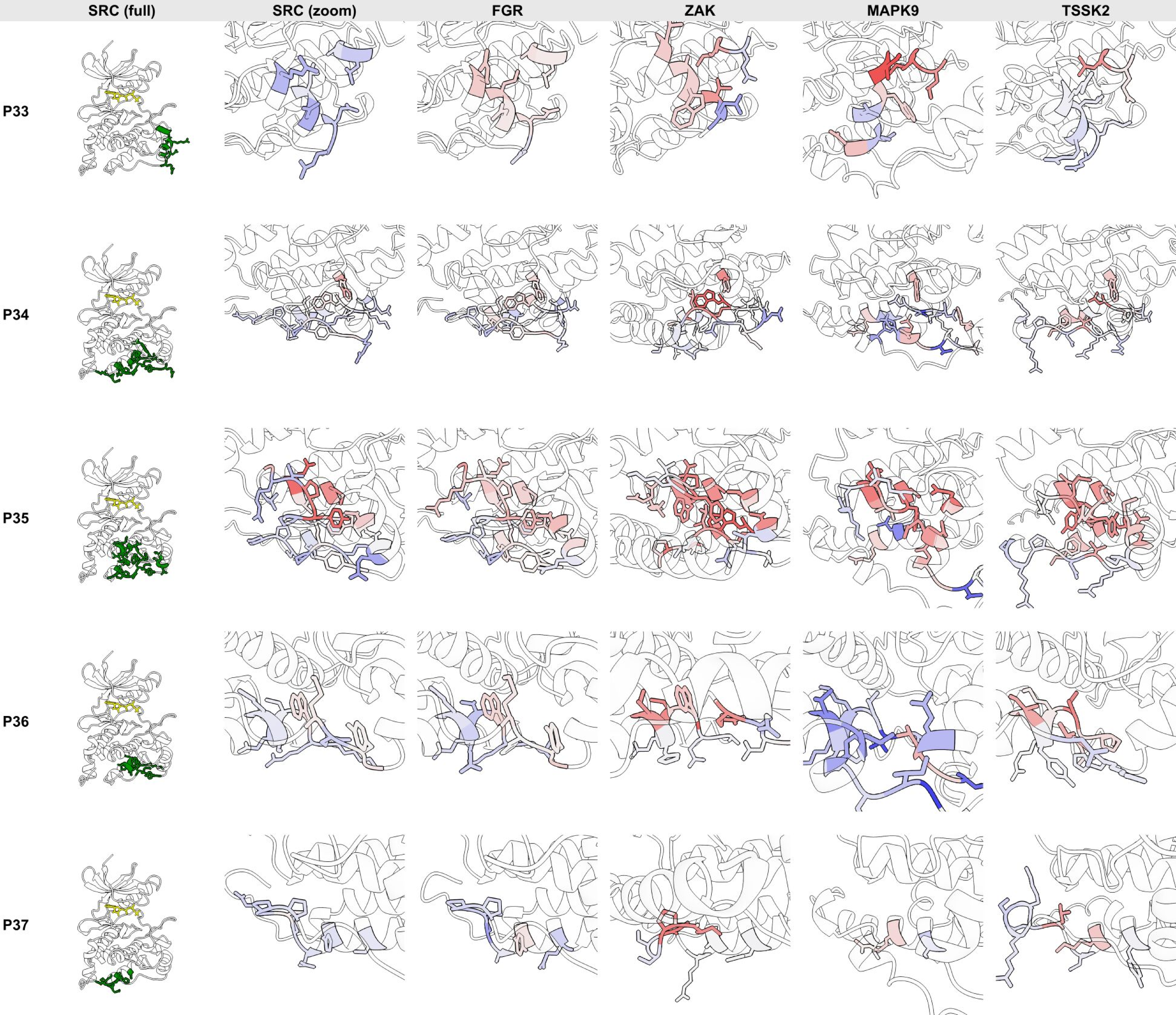
SRC (full)
SRC (zoom)
FGR
ZAK
MAPK9
TSSK2
P33
P34
P35
P36
P37
